# Molecular basis of proton-sensing by G protein-coupled receptors

**DOI:** 10.1101/2024.04.17.590000

**Authors:** Matthew K. Howard, Nicholas Hoppe, Xi-Ping Huang, Christian B. Macdonald, Eshan Mehrota, Patrick Rockefeller Grimes, Adam Zahm, Donovan D. Trinidad, Justin English, Willow Coyote-Maestas, Aashish Manglik

## Abstract

Three proton-sensing G protein-coupled receptors (GPCRs), GPR4, GPR65, and GPR68, respond to changes in extracellular pH to regulate diverse physiology and are implicated in a wide range of diseases. A central challenge in determining how protons activate these receptors is identifying the set of residues that bind protons. Here, we determine structures of each receptor to understand the spatial arrangement of putative proton sensing residues in the active state. With a newly developed deep mutational scanning approach, we determined the functional importance of every residue in proton activation for GPR68 by generating ∼9,500 mutants and measuring effects on signaling and surface expression. This unbiased screen revealed that, unlike other proton-sensitive cell surface channels and receptors, no single site is critical for proton recognition in GPR68. Instead, a network of titratable residues extend from the extracellular surface to the transmembrane region and converge on canonical class A GPCR activation motifs to activate proton-sensing GPCRs. More broadly, our approach integrating structure and unbiased functional interrogation defines a new framework for understanding the rich complexity of GPCR signaling.

**One-sentence summary:** The protonation networks governing activation of human pH-sensing GPCRs are uncovered by integrative cryo-EM and deep mutational scanning.

## Introduction

Homeostatic control of acid-base balance is vital for cellular and tissue physiology. Precise sensing of pH is fundamentally important to acid-base homeostasis. In humans and other animals, diverse cell surface proteins respond to changes in extracellular pH by sensing protons. While the majority of cell surface proton sensors are ion channels^1–5^, three G protein-coupled receptors (GPCRs) respond to changes in extracellular pH: GPR4, GPR65, and GPR68^6,7^. These receptors are expressed in diverse cells that regulate central pH homeostasis^8^, pH sensing in the immune system^9–11^, and vascular responses to pH^12^. Understanding and precisely manipulating the function of these proton sensing GPCRs holds promise for a range of diseases like inflammatory bowel disease^10,13^, osteoarthritis^14^, and certain cancers^15–17^.

Given the relevance of proton sensing GPCRs to pH-dependent physiology, it is important to understand how these receptors work at the molecular level. For pH sensing ion channels and transporters, a defined cluster of polar and charged residues is often ascribed as the proton recognition site^1–8,10,18^. This view, however, has remained controversial because it is often challenging to completely abolish proton sensitivity with targeted mutagenesis^19,20^. Several models have been proposed for proton recognition by proton sensing GPCRs. Proton-sensing GPCRs harbor an abundance of extracellular histidine residues that likely titrate at physiologically relevant pH levels; initial studies therefore ascribed these histidines as critical for proton sensing^7,21^. However, mutational studies suggest that histidines are dispensable for proton sensing - in GPR68, removal of all extracellular histidines does not abolish proton-driven activation^19,20^. A recent study employing parallel mutagenesis of titratable residues in GPR68 identified a conserved triad of buried acidic residues in proton-sensing GPCRs^20^. Here, too, neutral mutations to these sites shift pH_50_ but do not abolish proton-mediated receptor activation. How protons activate proton-sensing GPCRs remains poorly defined.

Structural biology methods have revealed fundamental insights into the molecular recognition of diverse GPCR stimuli ranging from light, ions, small molecules, peptides and large proteins^22–28^. However, knowing the structural location and context of residues in a 3-dimensional structure does not immediately inform function. This is particularly true for proton sensing receptors, where individual protons are not readily resolved by modern X-ray crystallography and cryogenic-electron microscopy (cryo-EM) approaches. An ideal alternative would be comprehensive data for how every single residue contributes to proton sensation. Unfortunately, conventional mutagenesis strategies do not scale to the dozens of protonatable residues within proton sensors and the multiple substitutions required to carefully dissect effects of local charge and hydrogen bonding networks.

Deep mutational scanning (DMS) has emerged as a powerful method to probe protein function^29^. In this approach, comprehensive mutagenesis is combined with a sequencing-based pooled assay to systematically measure how individual substitutions at every single position in a protein affect protein function. A key requirement for DMS is a robust phenotype that can be used to dissect function in a pooled cellular assay. When combined with mechanistic readouts, DMS has uncovered the molecular basis of protein function, folding, and allostery^30–33^. For GPCRs, previous DMS studies have uncovered important residue-level contributions to cell surface expression^34–36^ or, less commonly and done separately, to signaling^37^. Conventional mutagenesis studies, however, routinely highlight that GPCR mutations influence both cell surface expression (e.g. due to changes in synthesis, folding or trafficking) and cellular signaling (either via direct effects on stimulus recognition, allosteric communication or signaling effector coupling). To quantify how mutations influence signaling therefore requires a new approach that can untangle the contribution of mutation effects on surface expression vs. signaling.

Integrating structural biology with deep mutational scanning could provide a new approach to decipher GPCR function, and is ideally suited to understanding how protons activate proton-sensing GPCRs. Here, we developed this integrated approach by: 1) determining cryo-EM structures of all three human proton sensing GPCRs and 2) developing a new method for mechanistic dissection of GPCR function by deep mutational scanning. We devised a sensitive cellular assay for GPCR signaling that is capable of differentiating the effects of every possible mutation on receptor activation. A parallel deep mutational scan of cell surface expression yielded a multi-phenotypic view of each mutation, which resolves fundamental ambiguities in the effect of each mutation on receptor function. We applied this new approach to GPR68 to identify critical residues responsible for proton sensing and for allosteric activation of G protein signaling. Integrating structures with comprehensive functional data yielded a comprehensive structure-function model for how a stimulus activates a GPCR.

### Receptor chimeras reveal distributed proton sensing

We first investigated whether a conserved site confers proton sensitivity in proton-sensing GPCRs analogous to proposed models for proton-sensing channels and transporters^1,3,4,38–40^. In HEK293 cells, GPR4, GPR65, and GPR68 activate cAMP signaling with distinct sensitivity to protons, which is reflected in pH_50_ values of 8.0, 7.4, and 6.7, respectively (**Fig. 1A**). We reasoned that if a single site is responsible for proton sensing, we could find it by swapping segments of one pH sensing receptor for another and looking for concordant changes in pH_50_. We chose GPR4 and GPR68 for this chimeric receptor experiment as they have the highest sequence identity (44%) but the largest difference in pH_50_.

**Fig. 1:**
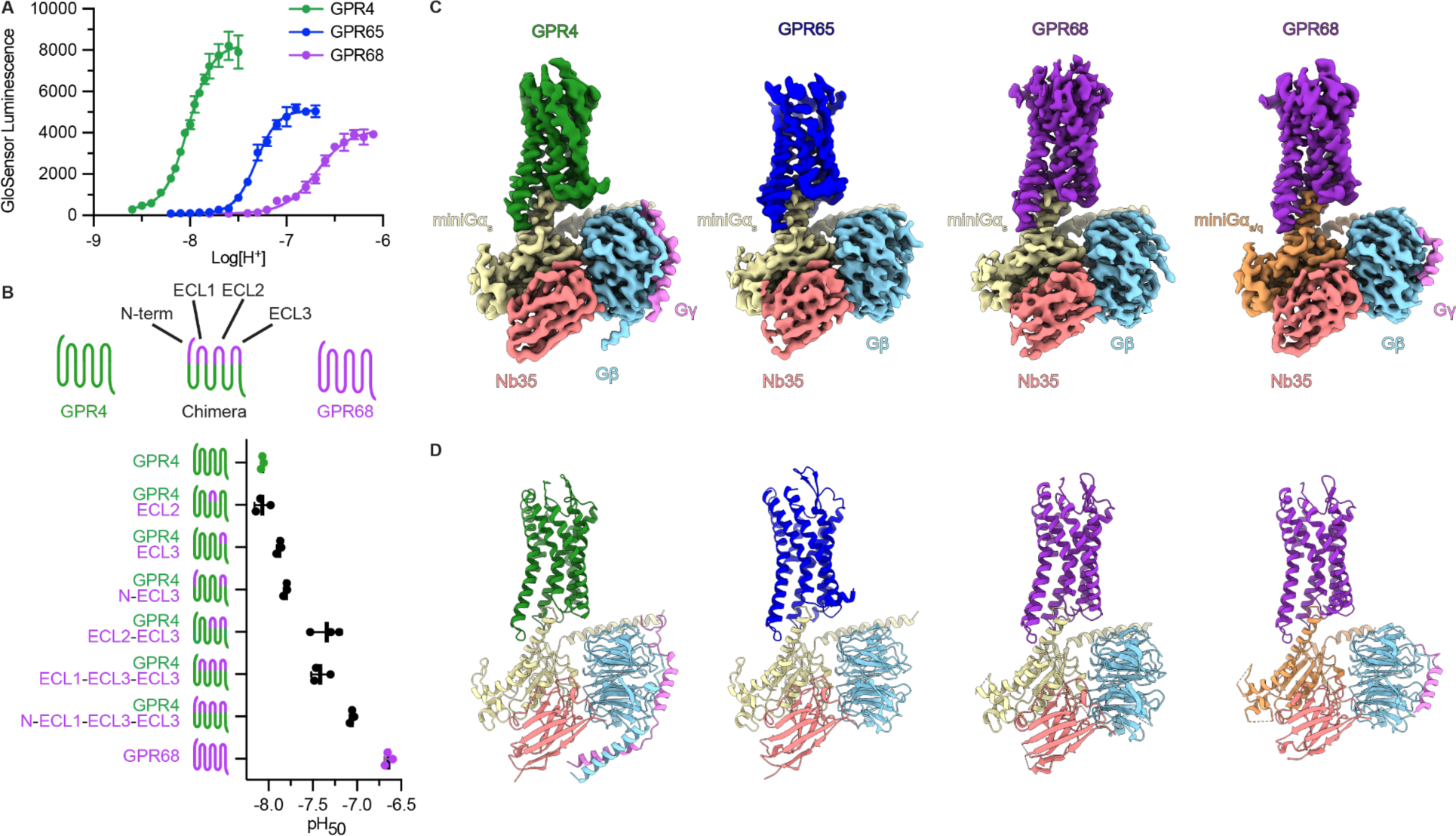
Chimeric pH sensors give insights into distributed proton sensing and Cryo-EM structures of GPR4, GPR65, and GPR68. (**A**) GloSensor cAMP accumulation assay showing the proton-sensing GPCRs, GPR4, GPR65, and GPR68, respond to decreasing pH. (**B**) GloSensor cAMP accumulation assay of GPR4-GPR68 chimeric receptors. Extracellular segments of GPR68 were grafted onto GPR4. A sequence alignment of GPR4 and GPR68 indicating swapped segments is available in **Fig. S1**. Data shown in **A, C** is from three independent biological replicates ± SD. (**C**) Cryo-EM density maps of GPR4-miniGα_s_, GPR65-miniGα_s_, GPR68-miniGα_s_, and GPR68-miniGα_s/q_. All four are bound to Gβγ and the stabilizing nanobody Nb35. (**D**) Ribbon model of each GPR4, GPR65, and GPR68 G protein complexes.

We designed chimeric receptors by grafting linear segments of the GPR68 extracellular regions onto GPR4 (**Fig. 1B, Fig. S1A**). Grafting points were chosen by matching the final Ballesteros-Weinstein (BW) position where GPR4 and GPR68 shared residue identity before diverging^41^ - this led to chimeric constructs that contain portions of GPR68 spanning extracellular loops (ECL) and the extracellular portions of the transmembrane (TM) helices. Each linear segment was tested individually and in combination with other segments in a cAMP accumulation assay (**Fig. S1B-E**). Out of the 15 constructs, 6 failed to show any proton-dependent signaling response, potentially because of deficits in folding or trafficking to the cell surface (**Fig. S1B-E**).

Chimeric constructs bearing single segments of GPR68 had little effect on pH_50_ (**Fig. 1B, Fig. S1B, Table S1)**. Introducing two segments of GPR68 into GPR4 also had little effect on pH_50_, with the exception of the ECL2/ECL3 chimera, which shifts the pH_50_ from 8.0 to 7.5 (**Fig. 1B, Fig. S1C, Table S1)**. Addition of the GPR68 ECL1 to this ECL2/ECL3 construct did not yield a further shift in pH_50_, although this construct is likely poorly expressed (**Fig. 1B, Fig. S1D, Table S1**). Paradoxically, addition of the GPR68 N-terminus to the ECL2/ECL3 construct restored pH_50_ to 8.0 (**Fig. S1D, Table S1)**. A final construct bearing the entire extracellular region of GPR68 grafted onto GPR4 yielded a pH_50_ of 7.1 (**Fig. 1B, Fig. S1E, Table S1)**.

These chimeric receptor experiments challenge a single site model of proton sensing in proton-sensing GPCRs. A single site of proton sensing would likely lead to a measurable shift in pH_50_ with exchange of a single segment. Instead, we find that substitution of individual extracellular segments of GPR68 is insufficient to cause a change in pH_50_ of the resulting chimera. Because swapping at least two segments yields a moderate shift in pH_50_, we conclude that a proton sensitive site in GPR68 is likely located in an interface between ECL2 and ECL3. Furthermore, because swapping the entire extracellular region of GPR68 is required for a pH_50_ that approaches that of native GPR68, we conclude that a network of proton sensitive sites is likely important for receptor activation.

### Cryo-EM structures of proton-sensing GPCRs

To understand how proton-sensing GPCRs recognize protons, we determined cryo-EM structures of human GPR4, GPR65, and GPR68 in complex with heterotrimeric G protein signaling subunits (**Fig. 1C, Fig. S2-5**). To overcome poor expression in HEK293 cells, we generated constructs of each proton-sensing GPCR fused C-terminally to miniGα proteins^42^. Both GPR4 and GPR65 have been previously characterized to drive cAMP production via activation of Gα_s_^7,21,42^; we therefore used miniGα_s_ to stabilize these receptors. By contrast, GPR68 has been shown to signal via Gα_q_ and Gα_s_^7,43^. We therefore used both miniGα_s_ and a chimeric miniGα_s/q_ construct to obtain structures of GPR68. We also screened different pH values for optimal high resolution reconstruction of receptor-G protein complexes. Although each receptor activates at distinct pH_50_ values when expressed heterologously in HEK293 cells, we found that purification at pH 6 enabled the best resolution for each receptor during single particle cryo-EM reconstruction. For GPR68-miniGα_s_, we included the positive allosteric modulator MS48107^19,44^, a derivative of ogerin^44^, in biochemical preparations. However, our structures did not reveal density for this ligand. Single particle reconstructions yielded nominal resolutions between 2.8-3.0 Å for the receptor-G protein complexes (**Table S2**). To improve reconstructions in the receptor extracellular regions, we also performed focused refinements on the 7TM domains (**Fig. S2-5**). The resulting maps enabled us to model each proton-sensing GPCR (**Fig. 1D**).

Structures of GPR4, GPR65, and GPR68 bound to miniGα_s_ revealed highly similar active-state conformations across the 7TM domains (RMSD < 1.5 Å) despite having sequence identities of 30-44% (**Fig 2A, Fig. S6**). Additionally, the conformation of GPR68 is highly similar between miniGα_s_ and miniGα_s/q_, with a RMSD of 0.9 Å (**Fig. S6**). Comparison to inactive and active structures of the prototypical class A GPCR, the β_2_-adrenergic receptor (β2AR) shows that each proton sensor is captured in a fully active conformation (RMSD of each proton sensor to active β2AR is <1.2 Å in the transmembrane regions) (**Fig. 2A**). This is reflected in a similar conformation of the common “P^5.50^I^3.40^F^6.44^” motif (superscripts indicate Ballesteros-Weinstein numbering^41^) (**Fig. 2C**); in the proton sensing GPCRs, a threonine and valine substitutes in the 3.40 position of GPR65 and GPR68, respectively, and valine is substituted for phenylalanine at the 6.44 position of GPR4 (**Fig. 2C**). In the proton sensors, the conserved GPCR “N^7.49^P^7.50^xxY^7.53^” motif harbors an aspartate at the 7.49 position, a substitution shared with ∼18% of all human Class A GPCRs (**Fig. 2D**). Finally, each proton receptor substitutes a phenylalanine in the conserved “C^6.47^W^6.48^xP^6.50^” motif in TM6. Each of these motifs adopts a similar conformation to active β2AR (**Fig. 2C-D**). While protons are a non-canonical stimulus, the activation pathway linking proton recognition to promotion of an active conformation is conserved between the proton sensing GPCRs and the broader class A GPCR family.

**Fig. 2:**
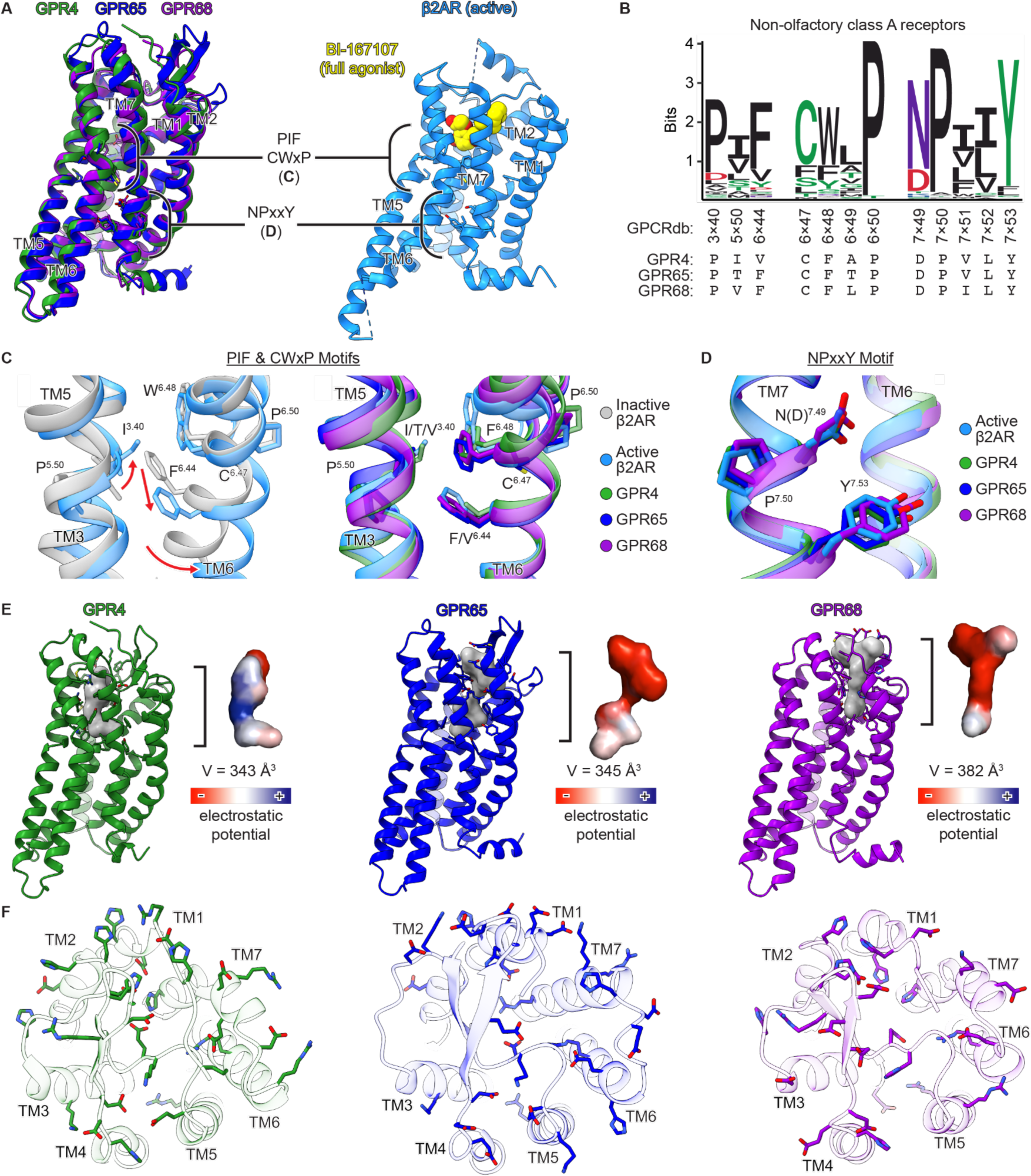
Structural features of human pH-sensing GPCRs. (**A**) (left) Structure alignment of 7TM domain GPR4-miniGα_s_, GPR65-miniGα_s_, and GPR68-miniGα_s/q_ shown as ribbons with sticks for indicated regions. (right) β2AR activated by full agonist BI-167107 (yellow) shown as ribbons with sticks for indicated regions (PDB ID: 3SN6). (**B**) Sequence Logo showing conservation of PIF, CWxP, and NPxxY among non-olfactory class A GPCRs. Sequences of GPR4, GPR65, and GPR68 are provided below for reference. (**C**) Close up views of the PIF and CWxP within the connector region in β2AR, GPR4, GPR65, and GPR68 show high similarity. Activation is associated with an outward movement of TM6 to accommodate G protein binding (PDB IDs: 2RH1 & 4LDO). (**D**) Close up views of the NPxxY motif in active-state GPR4, GPR65, GPR68, and β2AR activated by full agonist adrenaline (PDB ID: 4LDO). (**E**) Active-state models of GPR4, GPR65, and GPR68 each contain a charged pocket in the canonical orthosteric site. Each receptor is shown as a ribbon with residues lining each pocket shown as sticks. Pockets were calculated using CavitOmix, electrostatic surfaces were calculated using PyMol. (**F**) Each proton-sensing GPCR contains numerous titratable residues in the extracellular region. Extracellular regions of GPR4, GPR65, and GPR68 as viewed from the cell surface. Titratable residues are shown as sticks.

Structural diversity in the extracellular-facing domain of GPCRs enables recognition of a broad range of stimuli. We next compared similarities and distinctions in this region between the proton-sensing receptors and the broader Class A GPCR family. Each proton sensing receptor harbors an extracellular facing pocket that is lined by many polar and charged residues (**Fig. 2E**). Despite the presence of such cavities, our structures do not resolve density for potential activating ligands or metabolites that co-purify with the activated receptor. Given the size of these pockets, however, it is possible that endogenous metabolites or lipids may act as agonists or allosteric modulators for each of the three proton-sensing GPCRs. Indeed, the discovery of both positive and negative allosteric modulators for GPR4^45^, GPR65^46^, and GPR68^44,46^ supports the potential importance of this central cavity in modulating proton-sensing receptor function.

A unique feature of proton-sensing GPCRs is a large network of proton-titratable residues in the extracellular domain of the receptor (**Fig. 2E-F**). In addition to an abundance of histidine residues, each of the proton sensing GPCRs harbors additional acidic and basic residues that engage in an extended network of hydrogen bonding bridged by polar residues. Each receptor has a distinct network, although there are several structurally conserved positions harboring proton-titratable residues (**Fig. 2F**). Many of these surround what would be a canonical Class A GPCR orthosteric site. Collectively, these residues may coordinate protonation network(s) extending from the extracellular surface of each receptor that terminate at buried titratable residues^20,47,48^. Without inactive state structures or the ability to directly see protons, however, it is challenging to determine which of the many titratable residues is important for proton sensing. Nevertheless, these structures yield an understanding of the organization of putative proton-sensing residues in each receptor.

### Deep mutational scanning of GPR68 pH response

We next aimed to understand which of the numerous proton-titratable residues observed in structures of proton-sensing GPCRs are responsible for proton sensing and response. Conventional structure-function approaches to understand GPCR function use targeted mutagenesis combined with signaling studies to ascribe function to specific residues. Mutagenesis studies for understanding proton-sensitivity often require multiple substitutions, e.g. protonation mimicking mutations and charge reversals, to precisely define the effect of protonation at a specific site. The scale of mutagenesis experiments required for a comprehensive and unbiased profiling would not be imminently feasible with conventional approaches. We therefore turned to Deep Mutational Scanning (DMS), which is a high-throughput technique that enables sequence-to-function insight by profiling libraries of protein variants in a pooled format^29^.

We first devised a sensitive assay to enable DMS for GPCR signaling. Paramount to any successful DMS is a high throughput assay capable of discerning minute differences in phenotype. Prior work establishing DMS of GPCRs used the prototypical β2-adrenergic receptor (β2AR) to profile the effect of missense substitutions at every residue^37^. This DMS was enabled by a cAMP-dependent transcriptional readout for receptor activation, with each variant coupled to unique RNA barcodes that could be quantified by deep seqencing^37^. While pioneering for GPCR-DMS, there are several challenges of this approach including accumulation of signal at baseline, low signal-to-noise, and potential barcode clashing. To circumvent these limitations, we engineered a FACS-seq approach that reliably measures receptor activity. In this system, receptor activation of Gα_s_ triggers cAMP production that acts via a transcriptional reporter to produce eGFP (**Fig. 3A**). While this approach is similar to many assays that use transcriptional reporters of GPCR activity, we introduced several modifications to maximize the signal-to-noise of the FACS-seq assay. To provide maximal sensitivity to cAMP, we used a novel synthetic cAMP Response Element (CRE) sequence architecture recently discovered by massively parallel profiling of transcriptional response element architectures^49^. Because many GPCRs are basally active, a central challenge with transcriptional reporters of activity is low dynamic range; highly sensitive systems are often saturated by basal activity that occurs prior to activation of the receptor by a desired stimulus. We used two approaches to circumvent this issue. First, precise control of cell surface receptor expression with doxycycline induction enabled titration of receptor levels that maximize the dynamic range. Second, we fused eGFP to a dihydrofolate-reductase degron that is stabilized with the small molecule trimethoprim (TMP). In the absence of TMP, eGFP is constitutively degraded. Addition of TMP simultaneously with GPCR activation enables integration of the eGFP signal only in the presence of stimulus.

**Fig. 3:**
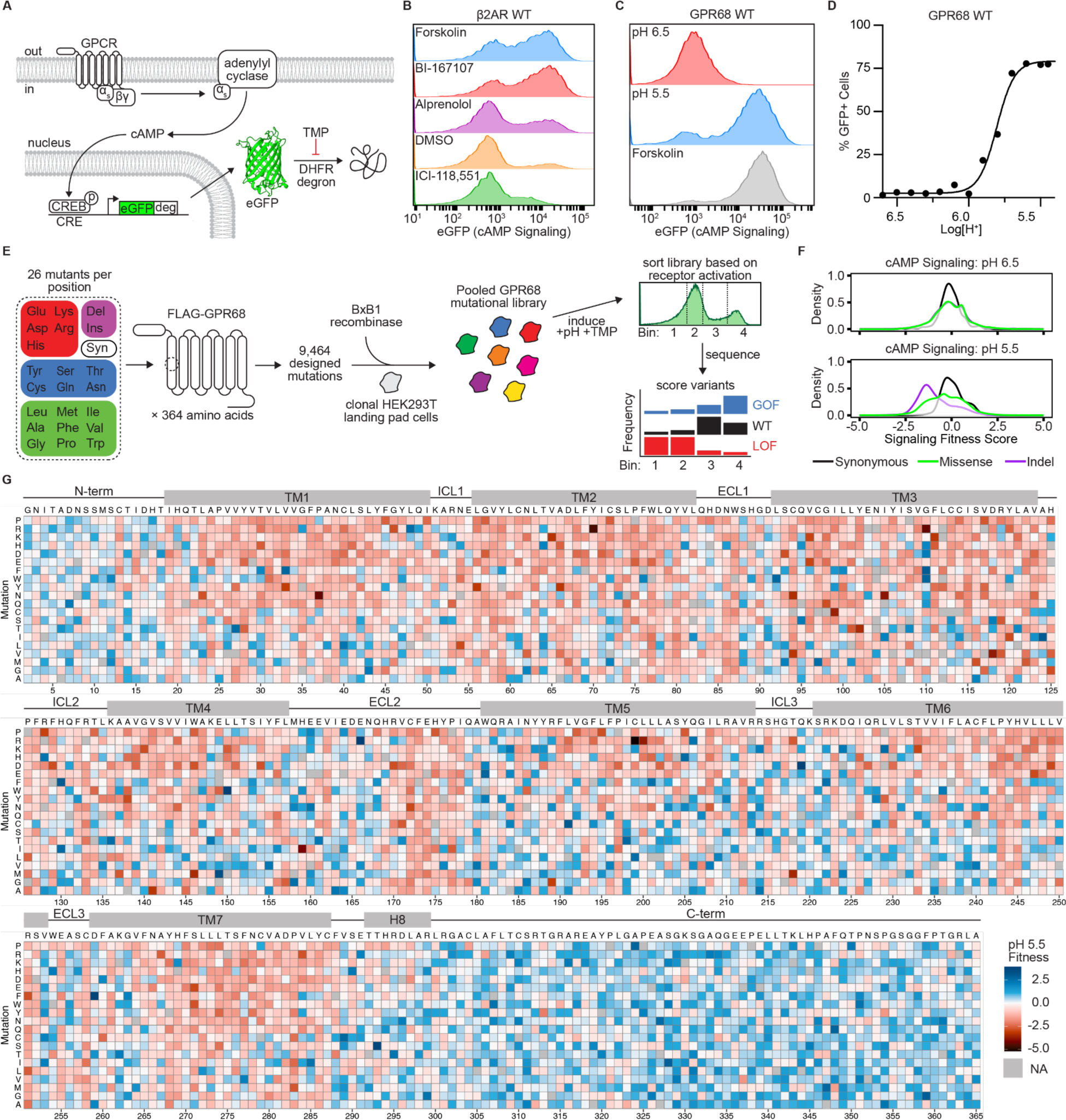
Deep mutational scan of GPR68 to determine critical residues for pH activation. (**A**) Schematic of cAMP transcriptional reporter assay. GPR68 activation triggers cAMP production leading to transcription of eGFP downstream of an engineered cAMP response element^49^. A dihydrofolate reductase (DHFR) degron eliminates background signal prior to stimulation. (**B**) Representative flow cytometry traces of β2AR treated with Forskolin (10 µM) which directly stimulates adenylyl cyclase, BI-167107 (10 µM, full agonist), Alprenolol (10 µM, antagonist), DMSO (0.1% v/v), and ICI-118,551 (10 µM, inverse agonist). (**C**) Representative flow cytometry traces of GPR68 at pH 6.5 (inactive) pH 5.5 (active), and at pH 7.5 with Forskolin (25 µM). (**D**) Representative pH dose-response curve for GPR68 WT. Arrows indicate pH condition shown in **C**. (**E**) Schematic of GPR68 mutational library generation and overview of FACS-seq pipeline for GPCR-DMS. (**F**) Distributions of variant effects of GPR68 signaling at pH 6.5 and pH 5.5. Fitness scores are relative to WT and were calculated using Enrich2.^53^ (**G**) Heatmap of cAMP signaling fitness scores for GPR68 mutational library at pH 5.5. WT sequence is shown above each section of heatmap, mutations are indicated on the left axis of each section, and the amino acid position is indicated by the numbers below each section. Positions and mutations with no data are shown as gray. Transmembrane helix cutoffs were determined using our GPR68 structure. Blue indicates increased cAMP signaling relative to WT, red indicates decreased cAMP signaling relative to WT. Data are fitness values from three biologically independent deep mutational scans.

We used β2AR to benchmark our assay and ensure that it provided adequate sensitivity and dynamic range. Using this system, we could reliably measure a full range of ligand efficacies. Forskolin treatment defined the ceiling of our assay, as it directly stimulated cAMP production via adenylyl cyclase (**Fig. 3B**). The full agonist BI-167107 produced a robust eGFP signal and closely mirrored the forskolin condition (**Fig. 3B**). A neutral antagonist, alprenolol, resulted in a modest eGFP signal over the DMSO vehicle baseline (**Fig. 3B**). This reflects alprenolol’s previously observed partial agonist activity at β2AR^50^. Further, an inverse agonist, ICI-118,551, demonstrated a reduction of eGFP signal relative to the DMSO treatment, concordant with its expected activity (**Fig. 3B**). These observations demonstrated that our system is capable of measuring differences in ligand-driven changes of Gα_s_-coupled receptor activation, and that it could similarly lend itself to measuring mutational effects on receptor activation in the context of DMS.

We next determined whether this transcriptional reporter assay can reliably detect pH dependent activation of proton-sensing GPCRs. Cell lines expressing GPR4 and GPR65 revealed significant basal signal at standard pH values required for cell culture (∼ pH 7.4). Attempts to increase the pH to decrease signaling were constrained by cell viability. By contrast, the eGFP signal for GPR68 is low at pH 7.4 and increased by approximately 30-fold upon addition of a pH stimulus, a change similar in magnitude to that induced by the direct adenylyl cyclase activator forskolin (**Fig. 3C-D**). The flow cytometry transcriptional reporter assay was performed in the absence of phosphodiesterase inhibitors and thus measures cAMP production instead of accumulation. In agreement with prior work, the pH_50_ of GPR68 is subsequently shifted ∼1 log unit to 5.8, compared to cAMP accumulation assays (**Fig. 1C, 3C**)^19,48^. We surmised that the transcriptional reporter provides a platform for DMS of GPR68.

To test mutational effects in an unbiased way using this reporter assay, we required a comprehensive mutational library of GPR68. Using the DIMPLE pipeline, we designed and generated a GPR68 DNA library containing all possible single missense mutations, a single synonymous mutation, as well as one, two, and three amino acid insertions and deletions at each position (**Fig. 3E**)^51^. This library of 9,464 variants was used to generate a pool of stable HEK293T cell lines where each cell contains only a single GPR68 variant thus enabling robust genotype-phenotype linkage (**Fig. 3E, Fig. S7**)^51,52^. With this pooled cell line library, we performed a screen at pH 5.5 (active) and pH 6.5 (inactive), as defined by the response of wild-type GPR68 (**Fig. 3C-D, Fig. S8-13**). To correlate phenotype and genotype, the pooled cell line library for each pH condition was sorted based on eGFP intensity into four bins using fluorescence-activated cell sorting (FACS) (**Fig. 3E**). The resulting subpopulations were sequenced, and a “fitness” score was calculated for each variant based on its distribution relative to synonymous mutations(**Fig. 3F**)^53^. These scores indicate whether a given variant is deleterious, beneficial or neutral for pH-dependent GPR68 activation and is plotted as a heatmap for the full length receptor at pH 5.5 (**Fig. 3G, Fig. S9-10**) and for pH 6.5 (**Fig. S11-12**).

Several features of this DMS provide confidence that this approach reliably measures the effect of mutations on GPR68 at scale. First, in the DMS at pH 6.5, mutations have very little effect on fitness scores (**Fig. 3F**). This is consistent with relatively little eGFP signal observed at the inactivating condition (**Fig. 3B, Fig. S8**). By contrast, at pH 5.5, we observe significant loss of fitness for regions of GPR68 likely important for function based on known structure-function relationships in the broader GPCR family (**Fig. 3F-G**). Specifically, the DMS fitness scores highlight that most substitutions in the transmembrane (TM) regions are poorly tolerated, while the amino and carboxy termini are less constrained (**Fig. S14A-B**). Additionally, substitution of cysteine residues known to form disulfide bonds between ECL2 and TM3 (residues 94 and 172) and the N-terminus and TM7 (residues 13 and 258) are universally deleterious (**Fig. S14B**). Substitutions to conserved GPCR motif positions as well as positions which interface with the G protein are also mostly deleterious (**Fig. S14B**). We concluded that the DMS of GPR68 activation using a transcriptional reporter of Gs signaling provides a comprehensive map of mutations and their effects.

### Integrative multi-phenotypic DMS

While DMS of GPR68 based on cAMP signaling activity provided initial insights into function, it is likely that many of the observed effects of mutations stem from changes in receptor surface expression. To deconvolve the effect of each mutant on surface expression vs. pH-dependent activation, we performed a second DMS based on surface expression of each GPR68 variant. Here we used fluorescently labeled anti-FLAG antibody to recognize an N-terminal FLAG tag on GPR68 (**Fig. 4A, Fig. S13, S15-16**). In this assay, surface expression is correlated with anti-FLAG signal; similar approaches are commonly used to measure the expression of single GPCR variants for structure-function studies. More broadly, we and others have used similar assays to surface expression of variant libraries of other membrane proteins^35,36,51,54–56^.

**Fig. 4:**
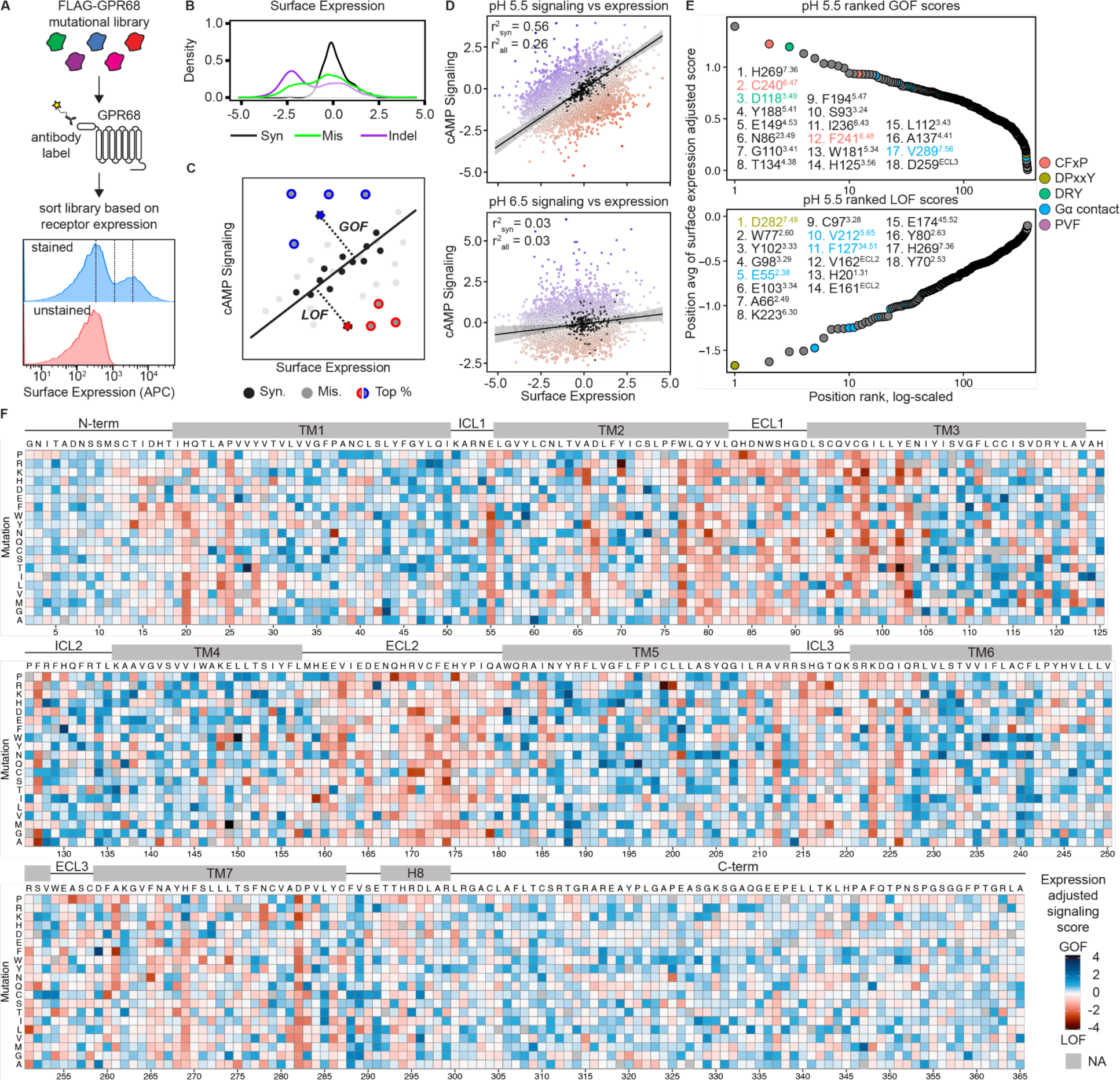
A multi-phenotypic screen reveals residues critical for GPR68 activation. (**A**) FLAG-GPR68 mutational library was labeled using an anti-FLAG antibody and receptor surface expression was measured using flow cytometry. Representative flow cytometry traces of stained (blue) and unstained (red) GPR68 mutational library. (**B**) Distribution of variant effects of GPR68 surface expression. Fitness scores are relative to WT. (**C**) Each mutation’s cAMP signaling score at each pH condition screened was plotted against its’ surface expression score. The euclidean distance of each mutant was calculated to a line fit to the population of synonymous mutations. Black points are synonymous variants, gray are missense variants, blue and red points are the top and bottom 2.5% missense mutants. (**D**) Scatter plot of surface expression vs cAMP signaling scores at pH 5.5 and pH 6.5. R^2^ values are shown for the synonymous (R^2^_syn_) and full missense (R^2^_all_) mutational library. (**E**) Surface expression-adjusted GOF and LOF pH 5.5 cAMP signaling scores are plotted in rank order. Positions are colored by sequence motif. Superscript corresponds to each residue’s Ballesteros-Weinstein number. (**F**) Heatmap of GPR68 mutational library surface expression-adjusted pH 5.5 cAMP signaling scores. WT sequence is shown above each section of heatmap, mutations are indicated on the left axis of each section, and the amino acid position is indicated by the numbers below each section. Positions and mutations with no data are shown as gray. Transmembrane helix cutoffs were determined using our GPR68 structure. Blue indicates higher activity relative to WT, red indicates lower activity relative to WT.

We next compared the effect of mutations on GPR68 activation and cell surface expression. As expected, synonymous mutations have little effect on GPR68 signaling or surface expression while insertions and deletions (Indels) have significant deleterious effects (**Fig. 3F, 4B**). Missense mutations are more distributed in effects for both surface expression and signaling (**Fig. 3F, 4B**). At pH 6.5, we see minimal effects of mutations because the receptor is inactive; rare missense mutations activate GPR68 (**Fig. 3F, Fig. S11).** To identify GPR68 mutations specifically important for pH-dependent activation, we calculated an expression-adjusted functional score for each variant. We first compare the effect of each mutation in the signaling and surface expression DMS; the resulting correlation indicates that activity in the signaling DMS is correlated to receptor surface expression (**Fig. 4C-D**). Synonymous mutations are expected to have minimal deleterious effects on expression or function - we use the correlation between signaling and expression scores of synonymous mutations to define a baseline regression fit for how expression levels influence signaling. We categorize mutations that have a higher than expected activity relative to their expression as gain-of-function (“GOF”). Conversely, mutations that have lower than expected function are loss-of-function (“LOF”). To identify GOF and LOF mutations, we calculated the euclidean distance of each missense mutation to the regression fit defined by synonymous mutations - missense substitutions with the most positive or negative scores yielded GOF and LOF mutations, respectively (**Fig. 4D, Fig. S17**).

Our analysis identifies the score for each individual substitution at a given GPR68 position. To identify individual sites with large effects on pH-dependent activation, we separated negative and positive distances scores and averaged for all missense substitutions at a given position. The resulting scores were then rank ordered, which provided a relative importance of each position for proton activation of GPR68 (**Fig. 4E**). This multi-phenotypic approach to integrating distinct effects of mutations enabled us to identify fundamental features of GPR68 activation (**Fig. 4E-F, Fig. S17**). Many positions that have substantial effects when mutated correspond to hallmark class A GPCR motifs such as the DRY, N(D)PxxY, CW(F)xP, and residues that contact the Gα protein (**Fig 4E-F, Fig. S17**)^57^. Intriguingly, numerous mutations identified in this screen correspond to ionizable residues in the extracellular regions of GPR68.

### Mechanism of GPR68 activation by protons

We next sought to integrate the cryo-EM structure of GPR68 with our expression-normalized DMS to build a structure-function map. We first reasoned that LOF mutations are likely to disrupt key interactions stabilized in the active conformation, and therefore visualized LOF scores for each residue position onto the active-state GPR68 cryo-EM structure (**Fig. 5A**). Conversely, we reasoned that GOF mutations are likely to disrupt interactions that stabilize inactive GPR68. Despite extensive attempts at obtaining a structure of inactive GPR68, we were unable to resolve this conformation of the receptor. We therefore use a model of GPR68 predicted by AlphaFold2 to be in an inactive-like conformation with an inward position of transmembrane helix 6 (**Fig. S18**). We visualized GOF scores for each residue using this AlphaFold model (**Fig. 5B**).

**Fig. 5:**
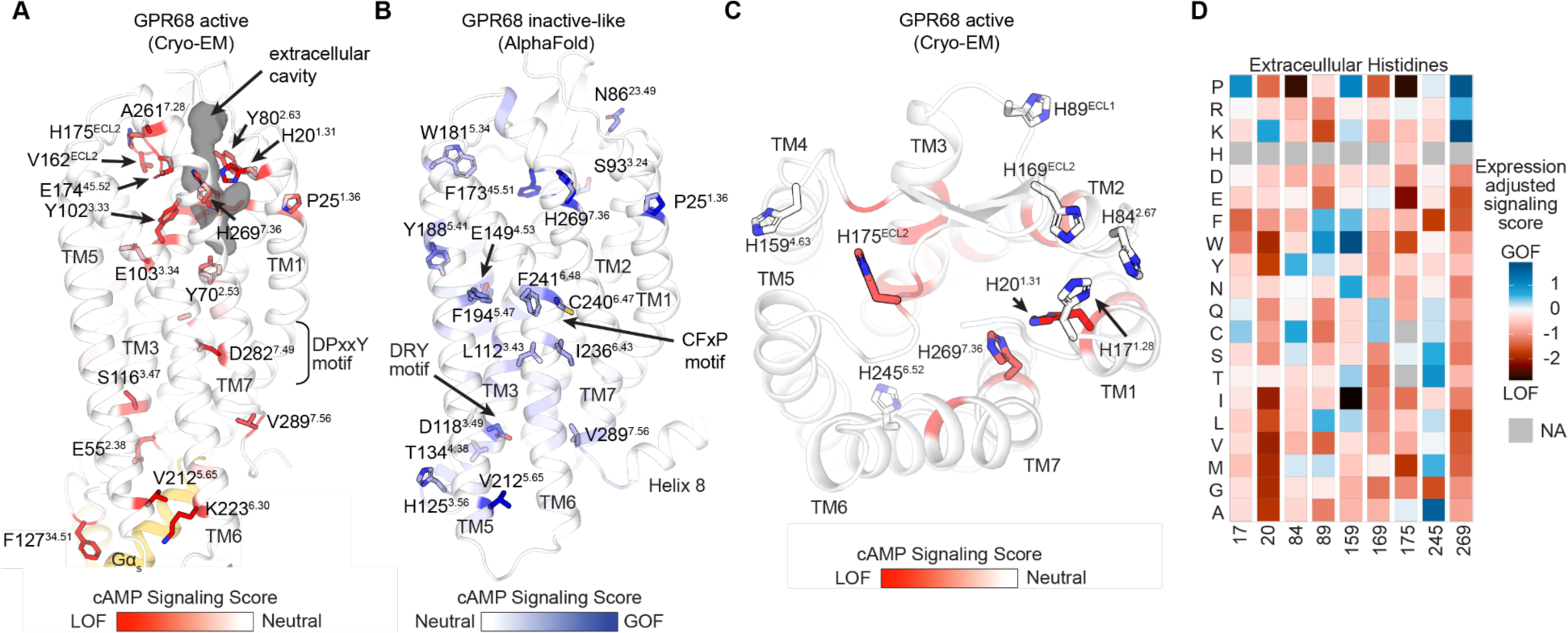
Structure mapping of GOF and LOF residues in GPR68 activation. (**A**) Residues where mutations result in increased cAMP signaling activity are shown as sticks on our experimental active state structure of GPR68. The extracellular cavity of GPR68 is shown as a grey surface. (**B**) Residues where mutations result in increased cAMP signaling activity are shown as sticks on an AlphaFold inactive-like structure of GPR68. Common class A GPCR activation motifs are indicated. (**C**) Top, extracellular, view of our active structure of GPR68 where all extracellular histidine residues are shown as stick and colored by their LOF score. (**D**) Subset heatmap from **4F** for each of the histidine residues shown in C.

Many GOF and LOF scoring mutations map to well-established class A GPCR motifs. For example, mutations in the D118^3.49^ in the DRY motif and C240^6.47^ and F241^6.48^ in the CW(F)xP motif lead to increased GPR68 signaling, consistent with important roles of these regions in stabilizing inactive GPCRs (**Fig. 5B, S18**)^57,58^. Mutations in D282^7.49^ in the N(D)PxxY motif lead to decreased signaling, supporting a key role of TM7 in receptor activation (**Fig. 5A, S18**). Additionally, mutation of F127^34.51^ in ICL2, which interacts directly with Gα, leads to a LOF (**Fig. 5A, S18**).

A more extensive set of LOF sites are adjacent to an extracellular facing cavity in GPR68 at a location similar to orthosteric sites in other class A GPCRs (**Fig. 5A**)^59,60^. Several LOF residues, including H269^7.36^, H20^1.31^, E174^45.52^, and Y102^3.33^ line this electronegative cavity, suggesting that this region is critically important for GPR68 activation by protons. We first looked more closely at histidine residues on the extracellular surface of GPR68 that have been proposed to be a critical determinant of proton-induced activation^19,47^. Our mutational scan provides an unbiased view on the relative importance of each histidine residue in GPR68 activity. Furthermore, the ability to test every amino acid substitution for function provides direct insight into how the charge state, hydrogen bonding interactions, and van der Waals interactions at a given position influence GPR68 activity. Two histidine residues in the extracellular region, H20^1.31^ and H269^7.36^, emerged as positions with a high LOF score in our global position analysis (**Fig. 5A,C-D and Fig. 6A-B**). Other histidines in the extracellular portion of GPR68 had more minor LOF effects (**Fig. 5C-D**). A closer analysis of substitutions in both H20^1.31^ and H269^7.36^ revealed that mutations to H269^7.36^ caused both gain and loss of function with positively charged substitutions leading to increased activity and negatively charged substitutions resulting in loss of activity (**Fig. 5D)**. For H20^1.31^, charge substitution leads to more subtle effects and the primary LOF score arises from hydrophobic substitutions (**Fig. 5D**). For both of these positions, we used a cAMP GloSensor assay to understand how substitution influences proton potency (**Fig. 6F**). For H269^7.36^, the amino acid sidechain pKa correlates with proton potency as indicated by the mutational scan. Intriguingly, the Hill slope of the proton response decreases from 4.16 in the wildtype receptor to 2.59-2.94 in the mutated receptors, suggesting that perturbing this position fundamentally alters cooperative proton binding to GPR68. For H20^1.31^, aspartate, asparagine, and arginine mutations led to subtle decreases in proton potency (**Fig. S18B**). Our mutagenesis screen therefore highlights that H269^7.36^ plays a central role in pH activation whereas histidines, like H20^1.31^, play secondary roles. More broadly, we observe large variability in the importance of each extracellular histidine and the effect of specific amino acid substitutions, suggesting that protonation of these other histidine residues is likely less important for GPR68 activation (**Fig. 5C-D**).

**Fig. 6:**
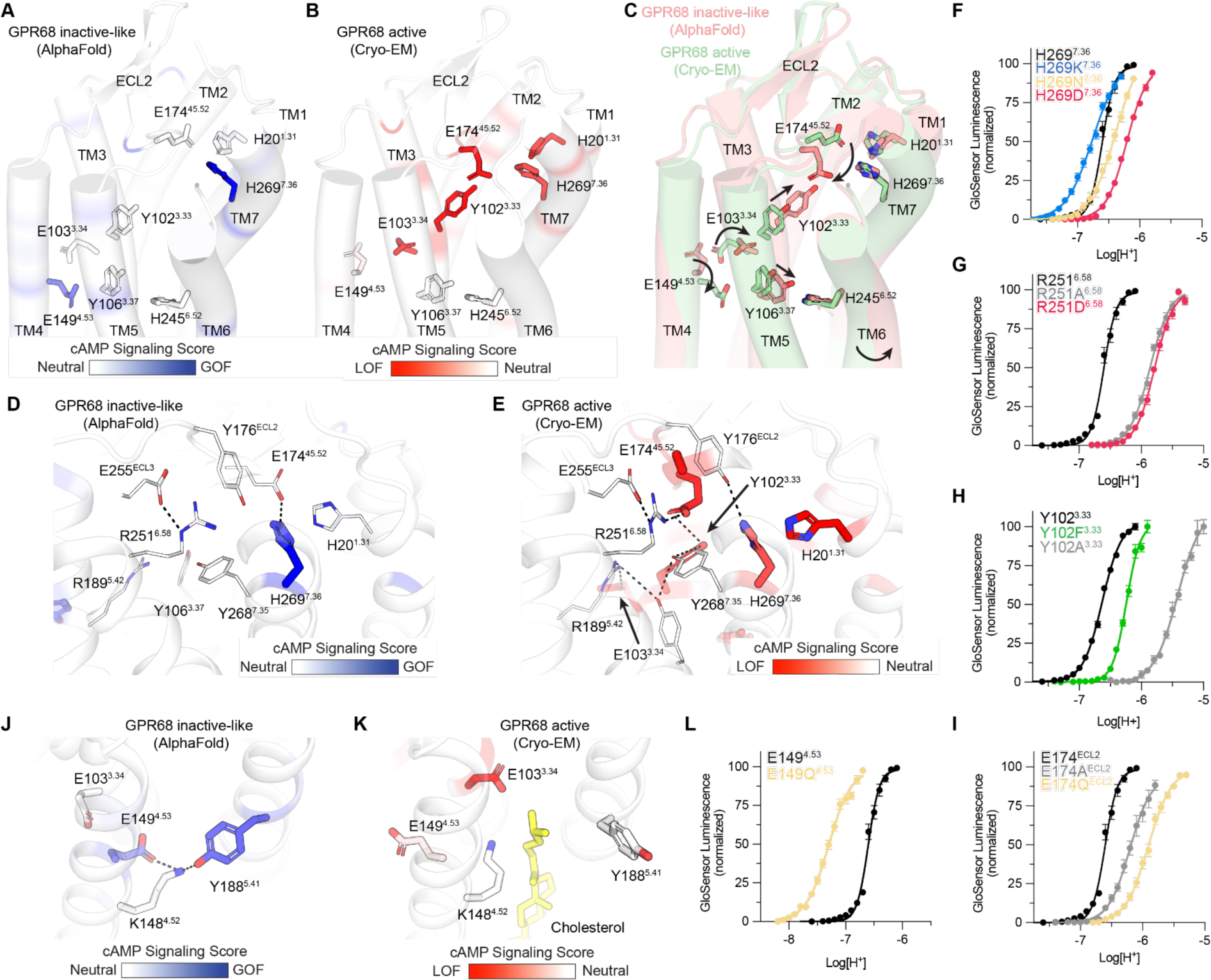
GPR68 activation network. (**A**) Mapping GOF positions onto the inactive-like GPR68 structure. (**B**) Mapping LOF positions onto the activated GPR68 Cryo-EM structure. Sticks are shows for key residues in activation network in **A** and **B**. (**C**) Overlay of inactive and active structures showing putative residue rearrangements upon proton activate of GPR68. (**D-E**) Key hydrogen bonds present in the active and inactive state networks. Residues are colored by their relative GOF or LOF score in each case. (**F-I**) cAMP accumulation GloSensor assays testing impact of mutations to key residues (**F**) H269, (**G**) R251, (**H**) Y102, and (**I**) E174. (**J-K**) Key hydrogen bonds present in the active and inactive state networks surrounding E149 and the cholesterol pocket. Residues are colored by their relative GOF or LOF score in each case. (**L**) cAMP accumulation GloSensor assays testing impact of neutral mutation to E149. Data shown in **F**-**I**, **L** is from three independent biological replicates ± SEM.

Using the DMS as a guide, we identified a network of interactions that connects the extracellular facing cavity to the core of the receptor (**Fig. 6A-B**). These interactions are predicted to rearrange when comparing the inactive-like conformation predicted by AlphaFold and our active-state cryo-EM structure of GPR68 (**Fig. 6C**). An extensive set of ionic and hydrogen-bonding interactions in active GPR68 engage LOF residues H269^7.36^ and E174^45.52^ (**Fig. 6E**). These interactions connect extracellular facing residues to two key residues in the core of GPR68: E103^3.34^ and E149^4.53^, which have strong LOF and GOF scores, respectively (**Fig. 6A-B**). Intriguingly, the extensive network of interactions in active GPR68 is rearranged in the AlphaFold predicted inactive-like state of GPR68 (**Fig. 6D**). Several conformational changes are notable.

First, activation of GPR68 is associated with a movement of E174^45.52^, which engages Y102^3.33^ and R251^6.58^ in the active state. Mutation of E174^45.52^, Y102^3.33^, or R251^6.58^ leads to a significant decrease in proton potency (**Fig. 6G-I**), supporting the importance of this interaction to GPR68 activation. The E174^45.52^:Y102^3.33^ interaction is associated with a rotation and upward displacement of TM3 that is relayed to E103^3.34^ and E149^4.53^ (**Fig. 6J-K**). In the AlphaFold prediction on inactive-like GPR68, E149^4.53^ engages K148^4.52^ and Y188^5.41^ in an intramembrane ionic interaction. This interaction is disrupted in active GPR68 by rearrangement of TM3 and the presence of the isooctyl chain of cholesterol that inserts between TM4 and TM5. This conformational rearrangement is supported by DMS results, which reveal that mutations in both E149^4.53^ and Y188^5.41^ are GOF. Indeed, the E149Q mutation, which mimics the protonated state, is more easily activated by protons (**Fig. 6L**). Our results confirm a prior study that identified E149^4.53^ as a critical activity associated residue in GPR68^20^, but provide critical structural context for this observation.

The combination of DMS and structural analysis therefore reveals that protonation of key residues surrounding an extracellular facing cavity, e.g. H269^7.36^ leads to a series of conformational rearrangements in GPR68 with TM3 as a central conduit. This relay converges to the conserved connector region in class A GPCRs that coordinates a rearrangement of the transmembrane helices to allow G protein binding and activation (**Fig. 6C)**.

### Tuning pH sensitivity in the proton sensor family

We next turned to examine whether GPR4 and GPR65 sense protons through a similar network of residues as GPR68. If they do, we would expect that the same positions critical for GPR68 activation are also important for the activation of GPR4 and GPR65. Across the family, there are several structurally conserved positions with ionizable residues (**Fig. 2F, Fig. S19**).

We first investigated the role of a conserved acidic residue within ECL2 (E170 in GPR4 and D172 in GPR65). Similar to GPR68, alanine and neutralizing mutations to this position cause a pronounced decrease in the cooperativity (Hill slopes) at each receptor (**Fig. S19, Tables S4-5**). The effect on proton potency is diminished in GPR65 and negligible in GPR4. These differential effects highlight the role that this residue plays in receptor activity. At higher pH, ECL2 is likely stabilized by several other interactions. At lower pH (e.g., in GPR68), this residue becomes more critical for stabilizing that conformation and thus also has a large effect on proton potency when mutated.

Both GPR4 and GPR65 have a negatively charged extracellular-facing cavity similar to GPR68. Having learned the charge dependence of GPR68 H269^7.36^, which is positioned at the top of this cavity, we tested the homologous set of mutations for GPR4 (H269^7.36^) and GPR65 (R273^7.36^). Indeed, at this site, we see that negatively charged residues universally decrease proton potency (**Fig. S19, Tables S4-5**). Positively charged residues at this position in GPR4 and GPR65 have less pronounced effects, perhaps highlighting that these positions are already protonated and at more basic pH conditions.

Finally, we examined the conserved glutamate, E^4.53^, which is in the middle of TM4 and buried far from both the extracellular solvent and the intracellular G protein binding pocket in each receptor. This position potentially serves as a key link between the proton-sensing network and residues involved in canonical activation motifs^47^. We hypothesized that the effects we demonstrated in GPR68 may thus hold true in GPR4 and GPR65 as well. In agreement with previous work, we observe an increase in potency for each receptor upon mutation to glutamine: GPR4 E145Q^4.53^ increases pH_50_ by ∼0.25. GPR65 E142Q^4.53^ increases pH_50_ by ∼0.1 and GPR68 E149Q^4.53^ increases pH_50_ by nearly a full pH unit (**Fig. S18-19, Tables S4-5**)^20^.

With these studies, we conclude that each receptor in the family shares a common buried acidic residue at which protonation likely drives activation. Furthermore, each receptor has a similar sensing mechanism on the extracellular side of the receptor, but the exact identity of the residues comprising it differs slightly between them.

## Discussion

Our integrative structural and deep mutational scanning studies suggest a general model for how protons activate the proton sensing GPCRs. Using GPR68 as a prototype of the proton sensing GPCR subfamily, we find that a network of amino acids connects an extracellular facing cavity to a conserved charged residue buried in the transmembrane core of the receptor. Protonation likely drives conformational changes in ECL2, which further stabilizes movement of TM3 and a series of rearrangements that connect the extracellular facing cavity to E^4.53^, a residue uniquely conserved in the proton sensing GPCRs. While we identify specific amino acids that are likely protonated upon activation, it is likely that additional sites bind protons upon receptor activation.

Several observations support such a distributed network of proton sensing. First, for each proton sensing receptor, cAMP assays reveal a pH Hill slope >4, suggesting significant cooperativity in proton-dependent activation. Second, our chimeric receptor constructs between GPR4 and GPR68 suggest that multiple distributed regions within the extracellular portions of the receptors define the pH setpoint. Finally, our DMS experiments do not identify a single cluster for GPR68, but instead many distinct residues in the extracellular regions with functional consequences. Although there are nuances to the proton-sensing domain of each proton-sending receptor, these networks converge upon hallmark GPCR motifs which link ligand binding to a conformational change allowing G protein binding. This provides an activation pathway from extracellular proton binding to G protein activation, and it points towards a conserved model across the family where GPCR proton sensing is not localized to a single site.

The distributed model for proton sensing-based GPCR activation contrasts with structure-guided mechanisms proposed for other membrane protein proton sensors and transporters. For many of these membrane proteins, proton-driven activation has been ascribed to single or small subsets of amino acids^1–3,38–40,61^. By contrast, our deep mutational scanning approach highlights that many protonatable residues contribute to proton-dependent activation in GPR68. A similar distributed network is likely important for GPR4 and GPR65. Although dramatic charge reversing substitutions at critical proton-recognition sites alter the pH_50_ of receptor activation, they do not ablate proton sensitivity in each of the proton sensing GPCRs. We speculate that this distinction in mechanism of proton sensitivity between GPCRs and ion channels may reflect the distinct biology associated with these proton sensors, with more distributed proton sensing networks in the proton sensing GPCRs being more amenable to tuning pH sensitivity over evolution.

Our approach to analyze the functional consequence of each amino acid in GPR68 provides key advances in deep mutational scanning to understand GPCR function. The cAMP-driven transcriptional reporter assay used to interrogate GPR68 is directly transferable to a large number of GPCRs that modulate cAMP, either by stimulating or inhibiting adenylyl cyclase. An additional advance is an engineered system that only integrates cAMP-driven transcriptional output in the presence of the receptor stimulus; this overcomes fundamental challenges with basal signaling suppressing the signal-to-noise of transcriptional readouts of GPCR activation. Perhaps the most important advance we introduce here is accounting for surface expression while evaluating the effect of any mutation on cAMP production. In the absence of such normalization, many loss or gain of function mutations simply reflect changes in receptor biogenesis or trafficking to the cell surface. By developing a way to integrate mutational scanning for multiple phenotypes, we unambiguously identified residues critical for GPR68 activation by protons.

More broadly, our integration of structural biology and deep mutational scanning is likely to provide a new foundation for interrogation of the rich complexity of GPCR function. While the present study examines only two phenotypes, future work could incorporate robust assays for other aspects of GPCR function, including signaling through different G protein and β-arrestin pathways, receptor internalization, location dependent signaling, and receptor biogenesis. While the power of our approach is clear for receptors with stimuli that are invisible to conventional structural biology, these integrative approaches are broadly able to bridge insights gained from biochemical and structural studies with the significant complexity of GPCR function in the cellular context. We envision that integrative interrogation of GPCR structures will reveal determinants of orthosteric and allosteric ligand binding, novel allosteric sites, and regions of receptors important in engaging signal transducers and regulatory complexes. Together, these approaches will allow the development of more quantitative models of receptor function, enable further therapeutic development, and uncover novel receptor biology.

## Materials and Methods

### GloSensor cAMP assays

Proton-sensing GPCR Gs activation and cAMP production were determined using the GloSensor cAMP assay. The following method was adopted from a previously published procedure with modifications^44^. In detail, HEK293T cells were maintained and cotransfected with receptor DNA and GloSensor cAMP reporter plasmids in DMEM containing 10% FBS. Overnight transfected cells were plated in poly-l-lysine coated 384-well white clear-bottom plates in DMEM supplemented with 1% dialyzed fetal bovine serum (dFBS), about 15,000 cells in 40 μL per well, for a minimum of 6 h up to 24 h. Assay buffers were prepared in 1x Calcium- and Magnesium-free HBSS supplemented with different organic buffer agents for different pH ranges, 20 mM MES for pH 5.00–6.60, 20 mM HEPES for pH 6.70–8.20, and 20 mM TAPS for pH 8.30–8.60. pH was adjusted with KOH at room temperature. PDE inhibitor Ro 20-1724 at final 10 µM was added to working solutions just before the assays. To stimulate cells with desired pH solutions, cells were first removed of medium (gently shaking off) and stimulated with desired pH solutions (25 µl/well) supplemented with 2 mM luciferin. The cell plate was incubated at room temperature for 20 - 30 min before luminescence was counted. For stimulation solutions with pH below 6.0, cells (medium was not removed) first received 10 µl pH 7.4 assay buffer containing luciferin (final 2 mM) and Ro 20-1724 (final of 10 µM) for a minimum of 30 min. After luciferin loading, medium and luciferin solutions were removed; cells were then stimulated with desired pH solutions containing 2 mM luciferin and 10 µM Ro 20-1724 as above. The cell plate was incubated at room temperature for 20 - 30 min before counting. Data presented in Figures here has been normalized to % max response or fold of basal, pooled for analysis using the built-in 4 parameter logistic function in the GraphPad Prism V10. Full tables of pharmacologic parameters can be found in **Tables S1, S3-5**.

### GPR68 Deep Mutational Scan

#### GPR68 deep mutational scanning library generation

Our DIMPLE platform was used to generate the GPR68 deep mutational library^51^. Briefly, we designed the library to contain all missense mutations at each position in GPR68. We additionally included synonymous mutations and insertions and deletions of 1, 2, and 3 amino acids at each position. These mutations were encoded in oligos with flanking BsaI sites and then ordered as a SurePrint Oligonucleotide library (Agilent Technologies)(**Table S6**). This DNA was resuspended and the sublibrary fragments were amplified using PrimeStar GXL DNA polymerase and fragment-specific primers (**Table S7**). These reactions were subjected to PCR cleanup using Zymo Clean and Concentrate-5 kits. The cDNA sequence of GPR68 WT was synthesized by Twist Bioscience in their High Copy Number Kanamycin backbone, BsmBI and BsaI cutsites were removed. For each library fragment, this plasmid was amplified to add BsaI sites, gel purified, and the corresponding oligo sublibrary were assembled using BsaI-mediated Golden Gate assembly. These reactions were cleaned and transformed into MegaX DH10B cells and added to 30mL LB + Kanamycin and grown while shaking until they reached OD 0.6-0.7. DNA was isolated using a Zymo Zyppy Plasmid Miniprep kit. Each sublibrary was quantified using Invitrogen Qubit dsDNA HS assay kit and pooled in equimolar ratios. This pooled library was then assembled into our landing pad compatible cAMP reporter vector containing a GSGSGS-P2A-PuroR cassette for positive selection. The sequences of our empty cAMP transcriptional reporter plasmid and GPR68 WT plasmid are provided in **Table S8**.

### GPR68 DMS cell line generation

The HEK 293T LLP-iCasp9 cells used in this study were a gift from Doug Fowler (UW)^52^. Cell lines for GPR68 WT and the GPR68 mutational library were generated as follows. 1ug of DNA was cotransfected with 1ug BxB1 recombinase (pCAG-NLS-BxB1, Addgene #51271) using 3.75uL lipofectamine 3000 and 5uL P3000 reagent in 6 wells of a 6 well plate. For GPR68 WT, 2 wells were transfected and pooled following selection. For the GPR68 library, 18 wells were transfected in parallel. Cells were cultured in “D10” media (DMEM, 10% dialyzed FBS, 1% sodium pyruvate, and 1% penicillin/streptomycin) inside humidified incubators at 37C and 5% CO_2_ The landing pad in the cell line contains a Tet-on promoter upstream of the BxB1 recombination site and a split rapamycin analog inducible dimerizable Casp-9. Two days after transfection, we induce with doxycycline hyclate (2ug/mL) and treat with 10nM AP1903. Recombined cells have shifted the iCasp-9 cassette out of frame while unrecombined cells will express the cassette and upon treatment with AP1903 die from iCasp-9 induced apoptosis. Cells were selected for 2 days in AP1903 after which they were transitioned back to D10 supplemented with doxycycline. After two days of recovery, cells were transitioned to D10 supplemented with both doxycycline and puromycin to select for cells that have proper in-frame, full-length assemblies. Following puromycin selection for two days, cells were transitioned to D10 and expanded before freezing down or using in subsequent assays.

### Fluorescence activated cell sorting

For flow-based assays and cell sorting, frozen stocks of cells were thawed and allowed to recover for several days in D10 media. 48h prior to starting the experiment, cells were split into an appropriate sized dish such that they reach ∼75% confluency by the start of the sort. 36h prior to starting the assay, cells were induced with doxycycline hyclate (2ug/mL). Doxycycline was subsequently washed out after 24h and cells were maintained in D10 for the remaining 12h prior to sorting. For the pH and pH + 30uM ogerin conditions, the pH of D10 media was adjusted using HCl on the same day as the assay. The cAMP assay was run as follows: cells were swapped to D10 (at indicated pH) with trimethoprim for 8h. After this incubation, cells were detached using TrpyLE Express, washed, and resuspended in BD FACS buffer. The surface expression assay was run similarly, cells were simply detached using TrypLE after induction, stained with M2 FLAG APC-Surelight antibody (Abcam), washed 3x, and then kept covered on ice prior to sorting. Cell sorting was performed using a Cytoflex SRT. Briefly, cells were gated on FSC-A and SSC-A for HEK293T cells, then FSC-A and FSC-H for singlets.

For the cAMP assay, we assessed activity using eGFP on the FITC-A channel, and for surface expression assays, the APC-A channel. For the cAMP sorting experiments, the population was split into four roughly equal populations (% cells) based on the most active condition, pH 5.5 + 30uM Ogerin. These gates were maintained for all subsequent samples. For surface expression assays, the population was largely bimodal, and we gated using the peaks of each distribution and the intervening trough. For sorting experiments we aimed to collect cells equal or greater than 100x the expected number of variants in our library.

### Genomic DNA extraction and sequencing

Following cell sorting, genomic DNA (gDNA) was extracted from cells using Quick-DNA Zymo Microprep Plus kits. All resultant gDNA was used as template for PCR to generate amplicons of the target gene using cell_line_for_5 and P2A_cell_line_rev primers (**Table S7**). PCR reactions were then concentrated using Zymo DNA Clean and Concentrator-25 kits, mixed with NEB Purple Loading dye (6x, no SDS) and run on a 1% agarose 1x TBE gel. Target amplicons were excised and purified using Zymo Gel DNA Recovery kits. Amplicon DNA concentrations were then quantified using Invitrogen Qubit dsDNA HS assay kit.

Libraries were prepared for deep sequencing using the Illumina Nextera XT DNA Library prep kit. Libraries were indexed using the IDT for Nextera Unique Dual Indexes. Then, the lengths of indexed libraries were quantified using the Agilent TapeStation HS D5000 assay and concentrations were determined using Invitrogen Qubit dsDNA HS assay kit. Samples were normalized and pooled and then paired-end sequenced (SP) on a NovaSeq6000.

### Next generation sequencing data processing

Sequencing files were obtained from the sequencing core as fastq.gz after demultiplexing. The experiment was processed using a DMS-specific pipeline we have developed^63^. The pipeline implemented the following steps: first, adapter sequences and contaminants were removed using BBDuk, then paired reads were error corrected with BBMerge and mapped to the reference sequence using BBMap with 15-mers (all from BBTools^64^). Variants in the mapped SAM file were called using the AnalyzeSaturationMutagenesis tool in GATK v4^65^. The output of this tool is a CSV containing the genotype of each distinct variant as well as the total number of reads for each sample. This was then further processed using a python script which filtered out sequences that were not part of the designed variants and then formatted input files for Enrich2^53^. Enrichment scores were calculated from the collected processed files using weighted least squares and normalized using wild-type sequences. The final scores were then processed and plotted using R. A copy of this processing pipeline, sequencing counts, and fitness scores has been deposited in the Github repositories listed in the data availability section.

### GPR68 Deep Mutational scanning data analysis

Deep mutational scanning data were analyzed in R as described in the text. All scripts used to make figures have been deposited in a Github repository listed in the data availability section.

### GPR4, GPR65, GPR68 purification and structure determination

#### Expression and purification of proton sensor active-state complexes

The human GPR4, GPR65, and GPR68 genes with an N-terminal influenza hemagglutinin signal sequence and Flag epitope tag were cloned into a pcDNA3.1/Zeo vector containing a tetracycline inducible cassette. The miniG proteins (miniG_s399_ for GPR4 and GPR65 and miniG_s/q70_ for GPR68) were fused to the C terminus of each proton sensor preceded by a glycine/serine linker and rhinovirus 3C protease recognition site^42^. The resulting fusion constructs were transfected into inducible Expi293F-TetR cells (Thermo Fisher) using the ExpiFectamine transfection reagent per manufacturer instructions. After 18 h, protein expression was induced with 1 µg/mL doxycycline hyclate for 24 h before collection by centrifugation. Pelleted cells were washed with 50 mL phosphate buffered saline, pH 7.5 before storage at −80 °C. For receptor purification, frozen cells were hypotonically lysed in 20 mM MES, pH 6, 1 mM EDTA, 160 µg/mL benzamidine, 2 µg/mL leupeptin for 10 min at 25 °C. The membrane fraction was collected by centrifugation, and the fusion protein was extracted with 20 mM MES, pH 6, 300 mM NaCl, 1% (w/v) lauryl maltose neopentyl glycol (L-MNG, Anatrace), 0.1% (w/v) cholesteryl hemisuccinate (CHS, Steraloids), 2 mM MgCl_2_, 2 mM CaCl_2_, 160 µg/mL benzamidine, 2 µg/mL leupeptin with dounce homogenization and incubation with stirring for one hour at 4 °C. The soluble fraction was separated from the insoluble fraction by centrifugation and was incubated in batch for 1 h at 4 °C with homemade M1–Flag antibody-conjugated Sepharose beads. Sepharose resin was then washed extensively with 20 mM MES, pH 6, 150 mM NaCl, 0.1% (w/v) L-MNG, 0.01% (w/v) CHS, 2 mM MgCl_2_, 2 mM CaCl_2_ and then with 20 mM MES, pH 6, 150 mM NaCl, 0.0075% (w/v) L-MNG, 0.00075% (w/v) CHS, 2 mM MgCl_2_, 2 mM CaCl_2_ prior to elution with 20 mM MES, pH 6, 150 mM NaCl, 0.0075% (w/v) L-MNG, 0.00075% (w/v) CHS, 5 mM EDTA, 0.2 mg/mL Flag peptide. Eluted protein was concentrated in a 100 kDa MWCO Amicon spin concentrator, and injected onto a Superdex200 Increase 10/300GL (Cytiva) gel filtration column equilibrated in 20 mM MES, pH 6, 150 mM NaCl, 0.0075% (w/v) L-MNG, 0.0025% glyco-diosgenin (GDN, Anatrace), and 0.0005% CHS. Monodisperse fractions were complexed with G_β1γ2_ heterodimer and Nb35 at 2 molar excess overnight at 4°C. The next day, the heterotrimeric complex was concentrated with a 100 kDa MWCO spin concentrator and excess G_β1γ2_ and Nb35 was removed via size-exclusion chromatography, using a Superdex200 Increase 10/300 GL column (GE Healthcare) equilibrated in 20 mM MES pH 6, 150 mM NaCl, 0.00075% (w/v) L-MNG, 0.00025% (w/v) GDN, and 0.0001% CHS. Resulting heterotrimeric complex was concentrated with a 100 kDa MWCO spin concentrator for preparation of cryo-EM grids. For GPR68 structures with Co^2+^, 10 µM Co^2+^ was added to all buffers. For GPR68 structure at pH 7.5, 20 mM HEPES pH 7.5 was substituted for 20 mM MES pH 6.

### Expression and purification of G_β1γ2_

Human G_β1γ2_ heterodimer was expressed in *Trichoplusia ni* Hi5 insect cells (Expression Systems) using a single baculovirus generated in *Spodoptera frugiperda* Sf9 insect cells (Expression Systems). A bicistronic pVLDual construct contained the G_β1_ subunit with a N-terminal 6 × His tag, and an untagged human G_γ2_ subunit. For expression, Hi5 insect cells were transduced with baculovirus at a density of ∼3.0 × 10^6^ cells per mL, grown with 27 °C shaking at 130 rpm. 48 h post-transduction, cells were collected and washed in a hypotonic buffer containing 20 mM HEPES, pH 8.0, 5 mM β-mercaptoethanol (β-ME), and protease inhibitors (20 µg/mL leupeptin, 160 µg/mL benzamidine). The membrane fraction was then separated by centrifugation and solubilized with 20 mM HEPES pH 8.0, 100 mM sodium chloride, 1.0% sodium cholate, 0.05% dodecylmaltoside (Anatrace), and 5 mM β-mercaptoethanol (β-ME). Solubilized G_β1γ2_ heterodimer was then incubated with HisPur Ni-NTA resin (Thermo Scientific) in batch. Bound G_β1γ2_ heterodimer was washed extensively and detergent was slowly exchanged to 0.1% (w/v) lauryl maltose neopentyl glycol (L-MNG, Anatrace) and 0.01% CHS before elution with 20 mM HEPES pH 7.5, 100 mM NaCl, 0.1% L-MNG, 0.01% CHS, 270 mM imidazole, 1 mM dithiothreitol (DTT), and protease inhibitors. Eluted G_β1γ2_ heterodimer was pooled and rhinovirus 3C protease was added to cleave the N-terminal 6 × His tag during overnight dialysis in 20 mM HEPES pH 7.5, 100 mM NaCl, 0.02% L-MNG, 0.002% CHS, 1 mM DTT, and 10 mM imidazole. To remove uncleaved G_β1γ2_, dialysed material was incubated with HisPur Ni-NTA resin in batch. The unbound fraction was then incubated for 1 h at 4 °C with lambda phosphatase (New England Biolabs), calf intestinal phosphatase (New England Biolabs), and Antarctic phosphatase (New England Biolabs) for dephosphorylation. Final anion exchange chromatography was performed using a MonoQ 4.6/100 PE (Cytiva) column to purify only geranylgeranylated heterodimer. The resulting protein was pooled and dialysed overnight in 20 mM HEPES pH 7.5, 100 mM NaCl, 0.02% L-MNG, and 100 µM TCEP, and concentrated with a 3 kDa centrifugal concentrator to a final concentration of 162 µM. Glycerol was added to a final concentration of 20%, and the protein was flash frozen in liquid nitrogen and stored at −80 °C until further use.

### Expression and purification of Nb35

A pET-26b vector containing the Nb35 sequence with a carboxy-terminal Protein C affinity tag was transformed into BL21 Rosetta Escherichia coli cells (UC Berkeley QB3 MacroLab) and inoculated into 8 L of Terrific Broth supplemented with 0.1% glucose, 2 mM MgCl_2_, and 50 µg/mL kanamycin. Cells were induced with 400 µM IPTG at A600 of 0.6 and allowed to express at 20 °C for 21 h. Collected cells were incubated SET Buffer (200 mM Tris pH 8.0, 500 mM sucrose, 0.5 mM EDTA) in the presence of protease inhibitors (20 µg/mL leupeptin, 160 μg/mL benzamidine) and benzonase. To initiate hypotonic lysis, two volumes of deionized water were added to the cell mixture after 30 min of SET buffer mixing. Following lysis, NaCl was added to 150 mM, CaCl_2_ was added to 2 mM, and MgCl_2_ was added to 2 mM and lysate was centrifuged to remove the insoluble fraction. Supernatant was incubated with homemade anti-Protein C antibody-coupled Sepharose. Nb35 was eluted with 20 mM HEPES pH 7.5, 100 mM NaCl, and 2 mM CaCl_2_, 0.2 mg/mL protein C-peptide, and 5 mM EDTA pH 8.0, concentrated in a 10 kDa MWCO Amicon filter and injected over a Superdex S75 Increase 10/300 GL column (Cytiva) size-exclusion chromatography column equilibrated in 20 mM HEPES pH 7.5, 100 mM NaCl. Monodisperse Nb35 fractions were pooled, concentrated, and flash frozen in liquid nitrogen for storage at −80 °C until further use.

### Cryo-EM vitrification, data collection and processing GPR4-G_s_ pH 6 complex

The GPR4-G_s_ pH 6 complex was concentrated to 14 mg/mL supplemented with 0.05% CHAPS (Thermo Fisher) and 3 µL was applied onto a glow-discharged 300 mesh 1.2/1.3 gold grid covered in a holey gold film (UltrAufoil). Excess sample was removed with a blotting time of 4 s and a blotting force of 1 at 4 °C prior to plunge freezing into liquid ethane using a Vitrobot Mark IV (Thermo Fisher). A total of 9,018 movies were recorded with a K3 detector (Gatan) on a Titan Krios (Thermo Fisher) microscope operated at 300 keV with a BioQuantum post-column energy filter set to a zero-loss energy selection slit width set of 20 eV. Movies were recorded using dose-fractionated illumination at a nominal magnification of 86,000x (physical pixel size of 0.86 Å/pixel) and a defocus range of −1 to −2.1 µm for a total dose of 50.7 e^-^/Å^2^. Exposure areas were acquired with image shift collection using EPU (Thermo Fisher). Movies of the GPR4-G_s_ pH 6 complex were motion-corrected and dose-fractionated using UCSF MotionCor2^66^. Corrected micrographs were imported into cryoSPARC v3^67^. for CTF estimation via the Patch Estimation job. Micrographs with estimated CTF fit resolution > 5 Å were removed before further processing. Templates for particle picking were generated from the same complex reconstructed from a previous 200 keV imaging session. Particle picking templates were low-pass filtered to 20 Å and used to pick 8,608,607 particles. After picking, particles were extracted in a 288 pixel box and Fourier cropped to 48 pixels before 3D classification with alignment using a 20 Å low-pass filtered reconstruction and three random reconstructures generated from a prematurely truncated ab initio reconstruction job, called “garbage collectors,” with the Heterogeneous Refinement job type. Two rounds of Heterogeneous Refinement yielded 2,501,915 particles that were re-extracted in the same box size cropped to 72 pixels and classified in a third Heterogeneous Refinement job. The resulting 1,453,906 particles were re-extracted in the same box cropped to 144 pixels. A fourth round of Heterogeneous Refinement and 2D classification, yielded 878,077 particles that were extracted without cropping. A final round of Heterogeneous Refinement yielded 439,296 particles that were refined using the Non-Uniform Refinement job type giving the final full-particle map. Finally, local refinement using an inclusion mask covering the 7TM domain was performed, using poses/shift Gaussian priors with standard deviation of rotational and shift magnitudes limited to 3° and 2 Å, respectively.

### GPR65-Gs pH 6 complex

The GPR65-G_s_ pH 6 complex was concentrated to 11 mg/mL supplemented with 0.05% CHAPS (Thermo Fisher) and 3 µL was applied onto a glow-discharged 300 mesh 1.2/1.3 gold grid covered in a holey gold film (UltrAufoil). Excess sample was removed with a blotting time of 4 s and a blotting force of 1 at 4 °C prior to plunge freezing into liquid ethane using a Vitrobot Mark IV (Thermo Fisher). A total of 8,294 movies were recorded with a K3 detector (Gatan) on a Titan Krios (Thermo Fisher) microscope operated at 300 keV with a BioQuantum post-column energy filter set to a zero-loss energy selection slit width set of 20 eV. Movies were recorded using dose-fractionated illumination at a nominal magnification of 105,000x (physical pixel size of 0.81 Å/pixel) and a defocus range of −1 to −2.1 µm for a total dose of 46 e^-^/Å^2^. Exposure areas were acquired with image shift collection using SerialEM 3.8^68^. Movies of the GPR65-G_s_ pH 6 complex were motion-corrected and dose-fractionated using UCSF MotionCor2^66^.

Corrected micrographs were imported into cryoSPARC v3.1 for CTF estimation via the Patch Estimation job^67^. Micrographs with estimated CTF fit resolution > 5 Å were removed before further processing. Templates for particle picking were generated from the same complex reconstructed from a previous 200 keV imaging session. Particle picking templates were low-pass filtered to 20 Å and used to pick 8,673,428 particles. After picking, particles were extracted in a 288 pixel box and Fourier cropped to 48 pixels before 3D classification with alignment using a 20 Å low-pass filtered reconstruction and “garbage collectors” with the Heterogeneous Refinement job type. Two rounds of Heterogeneous Refinement yielded 2,588,765 particles that were re-extracted in the same box size cropped to 74 pixels and classified in two Heterogeneous Refinement jobs. The resulting 1,637,819 particles were re-extracted in the same box cropped to 150 pixels and further classified with two rounds of Heterogeneous Refinement and 2D classification. The resulting 1,055,443 particles were refined using the Non-Uniform Refinement job type. Particles were exported using csparc2star.py from the pyem script package, and a mask covering the 7TM domain of GPR65 was generated using the Segger tool in UCSF ChimeraX and the Volume Tools utility in cryoSPARC^69,70^. The particles and mask were imported into Relion v3.0 and classified in 3D without alignment through three separate iterations^71^. Particles comprising the three highest resolution classes were reimported into cryoSPARC for Non-Uniform Refinement. Finally, particles were exported into cisTEM for 7TM local refinements using the Manual Refinement job type and low-pass filtering outside of the mask^72^.

### GPR68-G_s/q_ pH 6 complex

The GPR68-G_q_ pH 6 complex was concentrated to 4 mg/mL and 3 µL was applied onto a glow-discharged 300 mesh 1.2/1.3 gold grid covered in a holey carbon film (Quantifoil). Excess sample was removed with a blotting time of 4 s and a blotting force of 1 at 4 °C prior to plunge freezing into liquid ethane using a Vitrobot Mark IV (Thermo Fisher). A total of 6,650 movies were recorded with a K3 detector (Gatan) on a Titan Krios (Thermo Fisher) microscope operated at 300 keV with a BioQuantum post-column energy filter set to a zero-loss energy selection slit width set of 20 eV. Movies were recorded using dose-fractionated illumination at a nominal magnification of 105,000x (physical pixel size of 0.855 Å/pixel) and a defocus range of −1 to −2.1 µm for a total dose of 50 e^-^/Å^2^. Exposure areas were acquired with image shift collection using EPU (Thermo Fisher). Movies of the GPR68-G_q_ pH 6 complex were motion-corrected and dose-fractionated using UCSF MotionCor2^66^. Corrected micrographs were imported into cryoSPARC v3.1 for CTF estimation via the Patch Estimation job^67^. Micrographs with estimated CTF fit resolution > 5 Å were removed before further processing. Templates for particle picking were generated from the same complex reconstructed from a previous 200 keV imaging session. Particle picking templates were low-pass filtered to 20 Å and used to pick 6,764,523 particles. After picking, particles were extracted in a 288 pixel box and Fourier cropped to 72 pixels before 3D classification with alignment using a 20 Å low-pass filtered reconstruction and “garbage collectors” with the Heterogeneous Refinement job type. Two rounds of Heterogeneous Refinement yielded 2,774,555 particles that were re-extracted in the same box size cropped to 192 pixels and classified in an additional Heterogeneous Refinement job. The resulting 1,144,750 particles were refined using the Non-Uniform Refinement job type. Particles were exported using csparc2star.py from the pyem script package, and a mask covering the 7TM domain of GPR68 was generated using the Segger tool in UCSF ChimeraX and the mask.py pyem script^70–72^. The particles and mask were imported into Relion v3.0 and classified in 3D without alignment^71^. Particles comprising the highest resolution class were reimported into cryoSPARC for Non-Uniform Refinement. Finally, particles were exported into cisTEM for 7TM local refinements using the Manual Refinement job type and low-pass filtering outside of the mask^72^.

### GPR68-G_s_ pH 6 complex

The GPR68-G_s_ pH 6 complex was concentrated to 4 mg/mL and 3 µL was applied onto a glow-discharged 300 mesh 1.2/1.3 gold grid covered in a holey carbon film (Quantifoil). Excess sample was removed with a blotting time of 4 s and a blotting force of 1 at 4 °C prior to plunge freezing into liquid ethane using a Vitrobot Mark IV (Thermo Fisher). A total of 6,812 movies were recorded with a K3 detector (Gatan) on a Titan Krios (Thermo Fisher) microscope operated at 300 keV with a BioQuantum post-column energy filter set to a zero-loss energy selection slit width set of 20 eV. Movies were recorded using dose-fractionated illumination at a nominal magnification of 105,000x (physical pixel size of 0.83 Å/pixel) and a defocus range of −1 to −2.1 µm for a total dose of 49 e^-^/Å^2^. Exposure areas were acquired with image shift collection using SerialEM 3.8^68^. Movies of the GPR68-G_s_ pH 6 complex were imported into cryoSPARC v3.1 for motion-correction, dose-fractionation, and CTF estimation^67^. Micrographs with estimated CTF fit resolution > 5 Å were removed before further processing. Templates for particle picking were generated from the same complex reconstructed from a previous 200 keV imaging session. Particle picking templates were low-pass filtered to 20 Å and used to pick 7,064,401 particles. After picking, particles were extracted in a 288 pixel box and Fourier cropped to 48 pixels before 3D classification with alignment using a 20 Å low-pass filtered reconstruction and “garbage collectors” with the Heterogeneous Refinement job type. Two rounds of Heterogeneous Refinement yielded 2,524,876 particles that were re-extracted in the same box size cropped to 144 pixels and classified in an Heterogeneous Refinement job. The resulting 804,228 particles were refined using the Non-Uniform Refinement job type. Particles were exported using csparc2star.py from the pyem script package, and a mask covering the 7TM domain of GPR68 was generated using the Segger tool in UCSF ChimeraX and the mask.py pyem script^69,70^. The particles and mask were imported into Relion v3.0 and classified in 3D without alignment. Particles comprising the highest resolution classes were reimported into cryoSPARC for Non-Uniform Refinement^71^. Finally, particles were exported into cisTEM for two local refinements using the Manual Refinement job type and low-pass filtering outside of masks^72^. In the first local refinement, the previous 7TM mask was used, and the second local refinement used a full-particle mask.

### Model building and refinement

Model building and refinement began with the Alphafold2 predicted structures as the starting models, which were fitted into the experimental cryoEM maps using UCSF ChimeraX^73^. The model was iteratively refined with real space refinement in Phenix and manually in Coot and Isolde^74–76^. The cholesteryl hemisuccinate model and rotamer library were generated with the PRODRG server, docked using Coot, and refined in Phenix and Isolde^77^. Final map-model validations were carried out using Molprobity and EMRinger in Phenix.

## Acknowledgements

We thank all members of the Coyote-Maestas and Manglik labs for their helpful feedback and discussion as we conducted this project.

## Author contributions

M.K.H., P.R.G., D.T., and W.C.M. generated and cloned the GPR68 deep mutational library. M.K.H designed and performed deep mutational scan experiments with input and assistance from N.H., A.Z., J.E., A.M., and W.C.M.. C.M. processed raw NGS sequencing data. M.K.H. analyzed deep mutational scanning datasets with input from N.H., A.M., and W.C.M. N.H. cloned, expressed, and biochemically optimized the purification of proton sensor constructs for structural studies. N.H. performed cryo-EM data collection, with help from cryo-EM facilities, and data processing. N.H., E.M., and A.M. built and refined models of the proton sensors. M.K.H., N.H., and X.H. generated receptor constructs, performed signaling studies, and analyzed the data. M.K.H. and N.H. prepared figures with input from W.C.M. and A.M.. M.K.H., N.H., W.C.M., and A.M., wrote the manuscript, with edits and approval from all authors. W.C.M. and A.M. supervised the overall project.

## Data and materials availability

Coordinates for the GPR4-G_s_, GPR65-G_s_, GPR68-G_s_ and GPR68-G_s/q_ complexes have been deposited in the RCSB Protein Data Bank under accession codes XXXX, XXXX, XXXX, and XXXX respectively. EM density maps for GPR4-G_s_, GPR65-G_s_, GPR68-G_s_ and GPR68-G_s/q_ complexes have been deposited in the Electron Microscopy Data Bank under accession codes XXXXX, XXXXX, XXXXX, and XXXXX, respectively.

Sequencing data from the GPR68 deep mutational scan have been deposited in the NCBI Sequence Read Archive under bioproject PRJNA1062987.

The pipeline used to process GPR68 deep mutational scan sequencing data along with raw variant counts and fitness scores has been deposited on Github (https://github.com/odcambc/GPR68_processing, https://github.com/odcambc/GPR68_DMS_QC). All scripts used to analyze data and prepare figures has been deposited on Github: https://github.com/Coyote-Maestas-Lab/GPR68_DMS.

