## Supplementary Information for "Molecular basis of proton-sensing by G protein-coupled receptors"

### Supplementary Materials

#### Fig. S1: Chimeric receptor strategy to discern proton sensing.

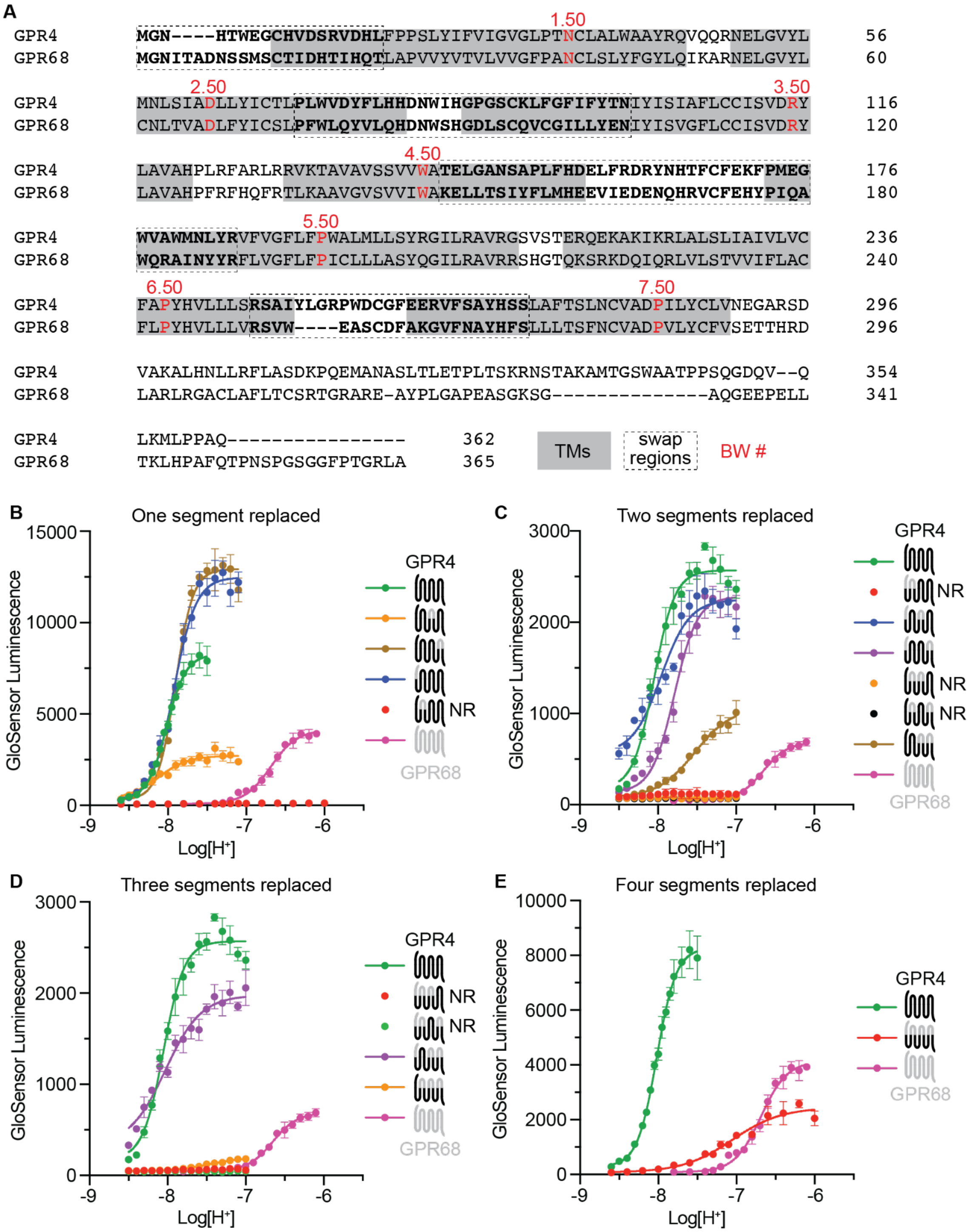


(**A**) Alignment of GPR4 and GPR68 generated using Clustal Omega. Grey indicates transmembrane helix regions. Dotted regions represent sequence segments that were grafted from GPR68 onto GPR4 to generate chimeric proton sensors.

(**B**-**E**) GloSensor cAMP accumulation assay showing proton activation for GPR4-GPR68 chimeras. All extracellular segments of GPR68 exchanged onto GPR4 are necessary to convert the proton response of GPR4 to be similar to the proton response of GPR68. Three or less segments is insufficient. Data are representative technical replicates from three independent biological replicates ± SD. Fits are shown for chimeras that retain proton sensitivity. Chimeras that do not demonstrate proton sensitivity are indicated by “NR” for no response, and no fit line is shown. A full table of pharmacologic parameters is available in **Table S1**.

**Fig. S2: Cryogenic electron microscopy processing of GPR4 miniG$_s$ pH 6.**

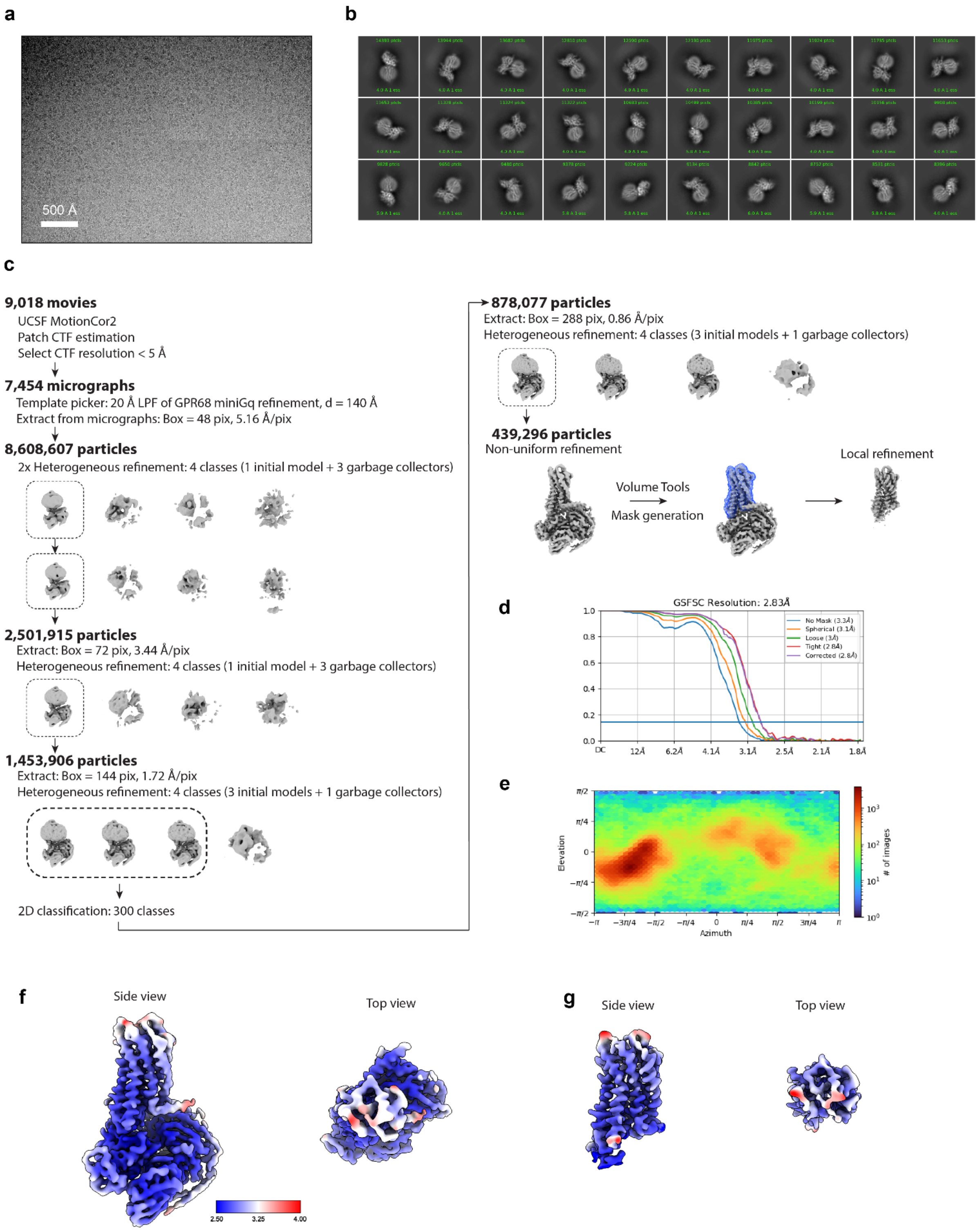


(**A**) A representative motion-corrected cryogenic electron microscopy (cryo-EM) micrograph obtained from a Titan Krios microscope**.** (**B**) A subset of highly populated, reference-free 2D-class averages. (**C**) Schematic showing the cryo-EM data processing workflow. Initial processing was performed using UCSF MotionCor2 and cryoSPARC. Particles were selected using iterative Heterogeneous refinement jobs followed by 2D classification. Finally, particles were processed using the local refinement job type with a 7TM mask. Dashed boxes indicated selected classes. (**D**) Gold-standard Fourier Shell Correlation (GSFSC) curve for final full-particle map computed in cryoSPARC. (**E**) Euler angle distribution of final full-particle map computed in cryoSPARC.(**F**) Side view and top view of local resolution for the final full-particle map of GPR4-$G_s$ pH 6 complex computed with local resolution in cryoSPARC. (**G**) Side view and top view of local resolution for the focused 7TM map of GPR4-$G_s$ pH 6 complex computed with local resolution in cryoSPARC.

**Fig. S3: Cryogenic electron microscopy processing of GPR65 miniG$_s$ pH 6.**

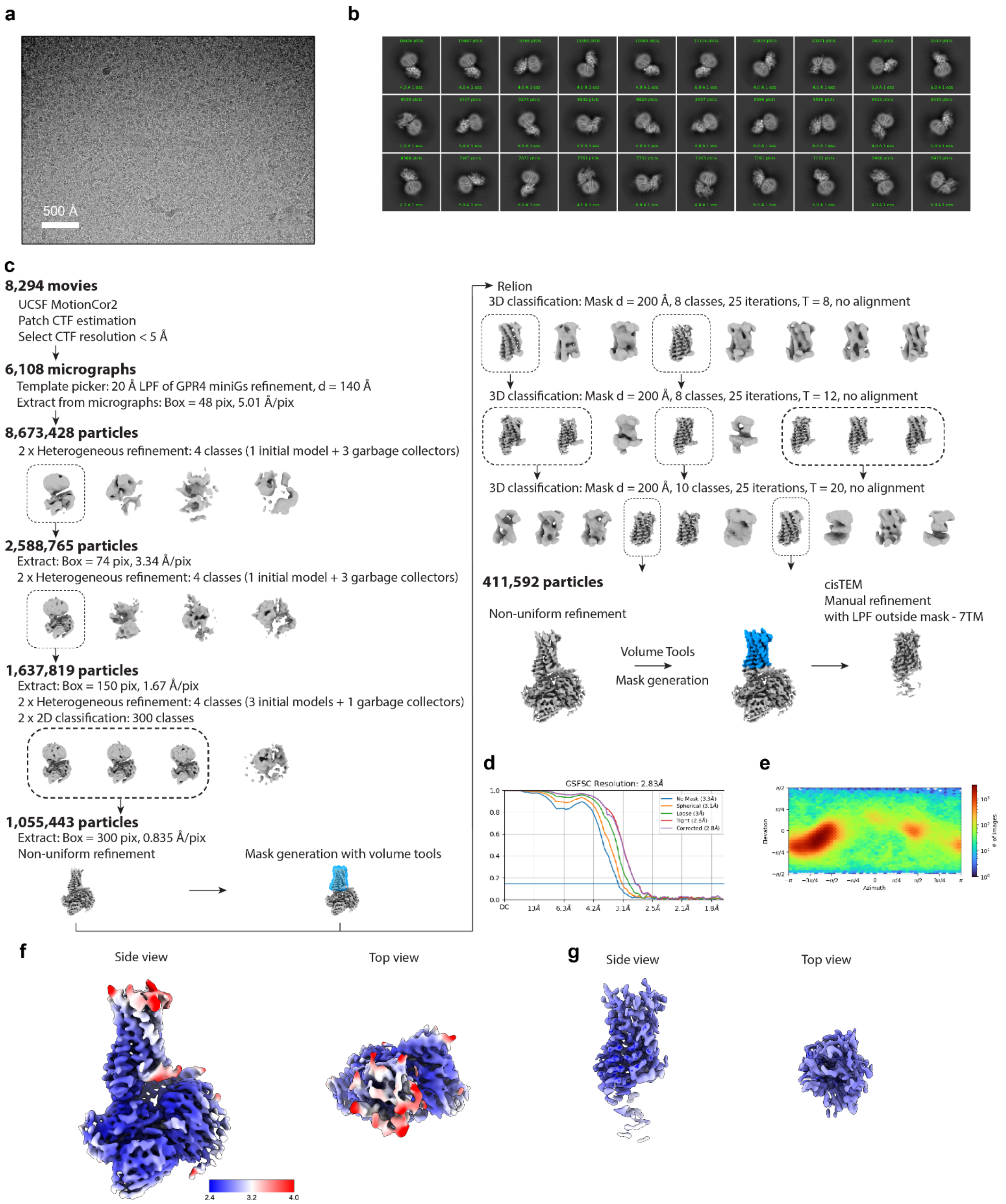


(**A**) A representative motion-corrected cryogenic electron microscopy (cryo-EM) micrograph obtained from a Titan Krios microscope. (**B**) A subset of highly populated, reference-free 2D-class averages. (**C**) Schematic showing the cryo-EM data processing workflow. Initial processing was performed using UCSF MotionCor2 and cryoSPARC.

Particles were transferred using the pyem script package to RELION for alignment-free 3D classification. Finally, particles were processed in cisTEM using the manual refinement job type with a 7TM mask followed by a full particle mask. Dashed boxes indicated selected classes. (**D**) Gold-standard Fourier Shell Correlation (GSFSC) curve for final full-particle map computed in cryoSPARC. (**E**) Euler angle distribution of final full-particle map computed in cryoSPARC. (**F**) Side view and top view of local resolution for the final full-particle map of GPR65-$G_s$ pH 6 complex computed with local resolution in cryoSPARC. (**G**) Side view and top view of local resolution for the focused 7TM map of GPR65-$G_s$ pH 6 complex computed with local resolution in cryoSPARC.

**Fig. S4: Cryogenic electron microscopy processing of GPR68 miniG$_{s/q}$ pH 6.**

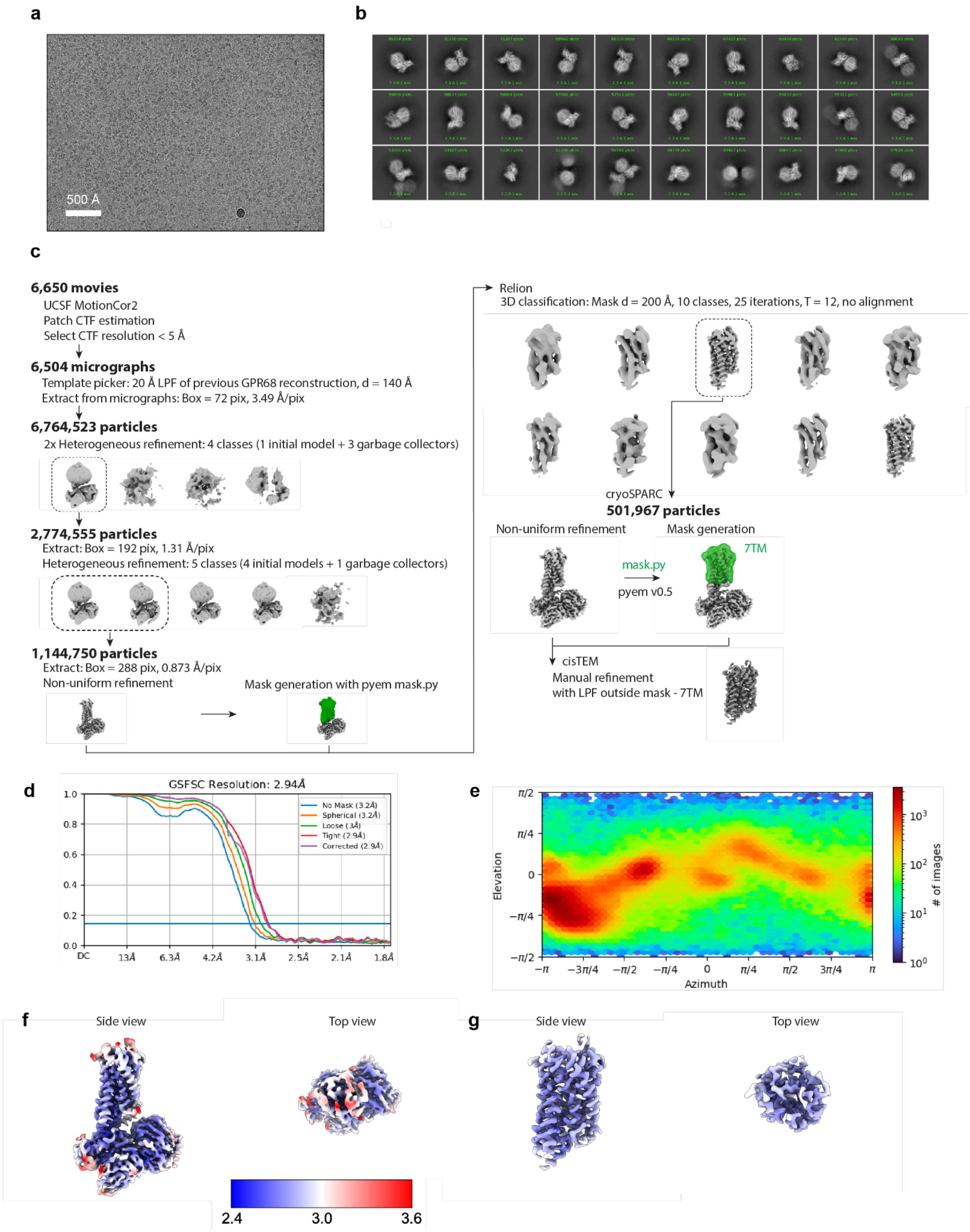


(**A**) A representative motion-corrected cryogenic electron microscopy (cryo-EM) micrograph obtained from a Titan Krios microscope. (**B**) A subset of highly populated, reference-free 2D-class averages. (**C**) Schematic showing the cryo-EM data processing workflow. Initial processing was performed using UCSF MotionCor2 and cryoSPARC. Particles were transferred using the pyem script package to RELION for alignment-free 3D classification. Finally, particles were processed in cisTEM using the manual refinement job type with a 7TM mask. Dashed boxes indicated selected classes. (**D**) Gold-standard Fourier Shell Correlation (GSFSC) curve for final full-particle map computed in cryoSPARC. (**E**) Euler angle distribution of final full-particle map computed in cryoSPARC. (**F**) Side view and top view of local resolution for the final full-particle map of GPR68-$G_q$ pH 6 complex computed with local resolution in cryoSPARC. (**G**) Side view and top view of local resolution for the focused 7TM map of GPR68-$G_q$ pH 6 complex computed with local resolution in cryoSPARC.

**Fig. S5: Cryogenic electron microscopy processing of GPR68 miniG$_s$ pH 6 with ms48017**

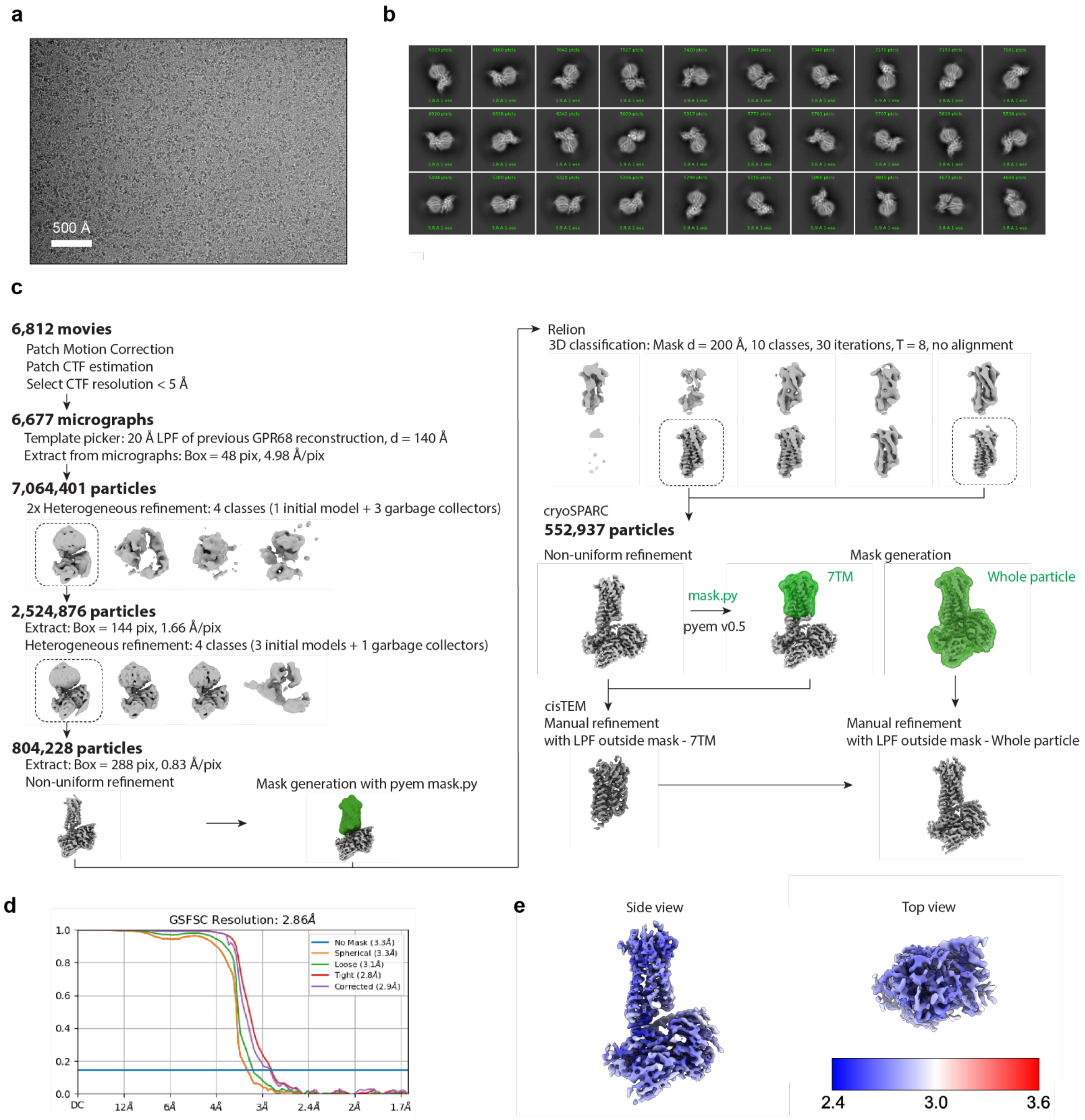


(**A**) A representative motion-corrected cryogenic electron microscopy (cryo-EM) micrograph obtained from a Titan Krios microscope**.** (**B**) A subset of highly populated, reference-free 2D-class averages. (**C**) Schematic showing the cryo-EM data processing workflow. Initial processing was performed using UCSF MotionCor2 and cryoSPARC. Particles were transferred using the pyem script package to RELION for alignment-free 3D classification. Finally, particles were processed in cisTEM using the manual refinement job type with a 7TM mask followed by a full particle mask. Dashed boxes indicated selected classes. (**d**) Gold-standard Fourier Shell Correlation (GSFSC) curve for final full-particle map computed in cryoSPARC. (**E**) Side view and top view of local resolution for the final full-particle map of GPR68-$G_s$ pH 6 complex computed with local resolution in cryoSPARC.

**Fig. S6: Proton sensor sequence identity and structural RMSD matrices**

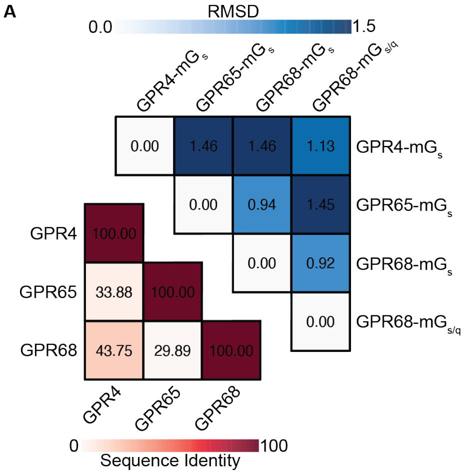


(**A**) GPR4, GPR65, and GPR68 sequence identity matrix and RMSD matris between our GPR4, GPR65, and GPR68 structures.

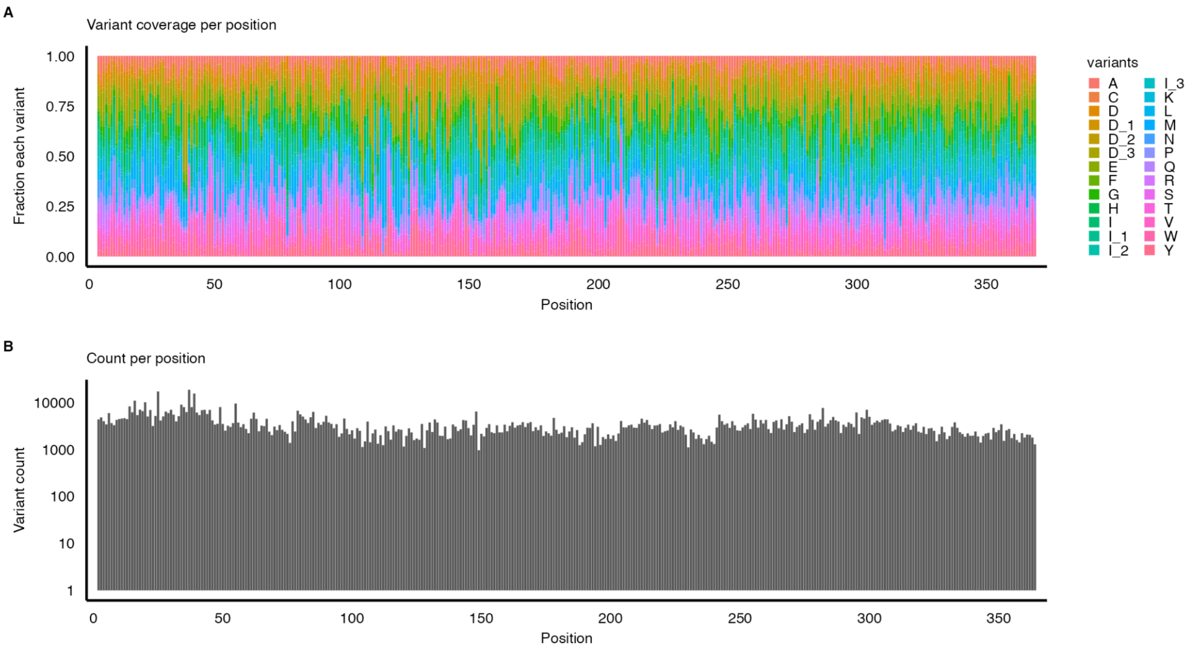


(**A**) Normalized counts of each variant per position in the GPR68 deep mutational library demonstrate relatively even overage of each possible mutation at each position. Variants are colored by their single letter amino acid abbreviation, or "I_#" and "D_#" for insertions of G, GS, GSG (# = 1-3) or deletions of 1, 2, or 3 amino acids. (**B**) Total counts per position in GPR68 deep mutational library demonstrates relatively even total coverage for each position.

**Fig. S8: Comparison of FACS eGFP signal for GPR68 mutational library cAMP signaling at pH 5.5 and pH 6.5**

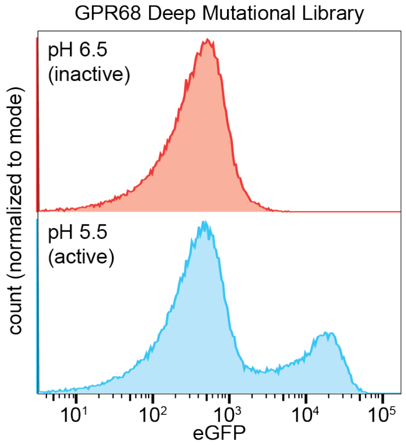


(**A**) Representative plot of GPR68 mutational library eGFP distribution at both pH 5.5 and pH 6.5.

**Fig. S9: GPR68 cAMP signaling at pH 5.5 DMS fitness score full heatmap**

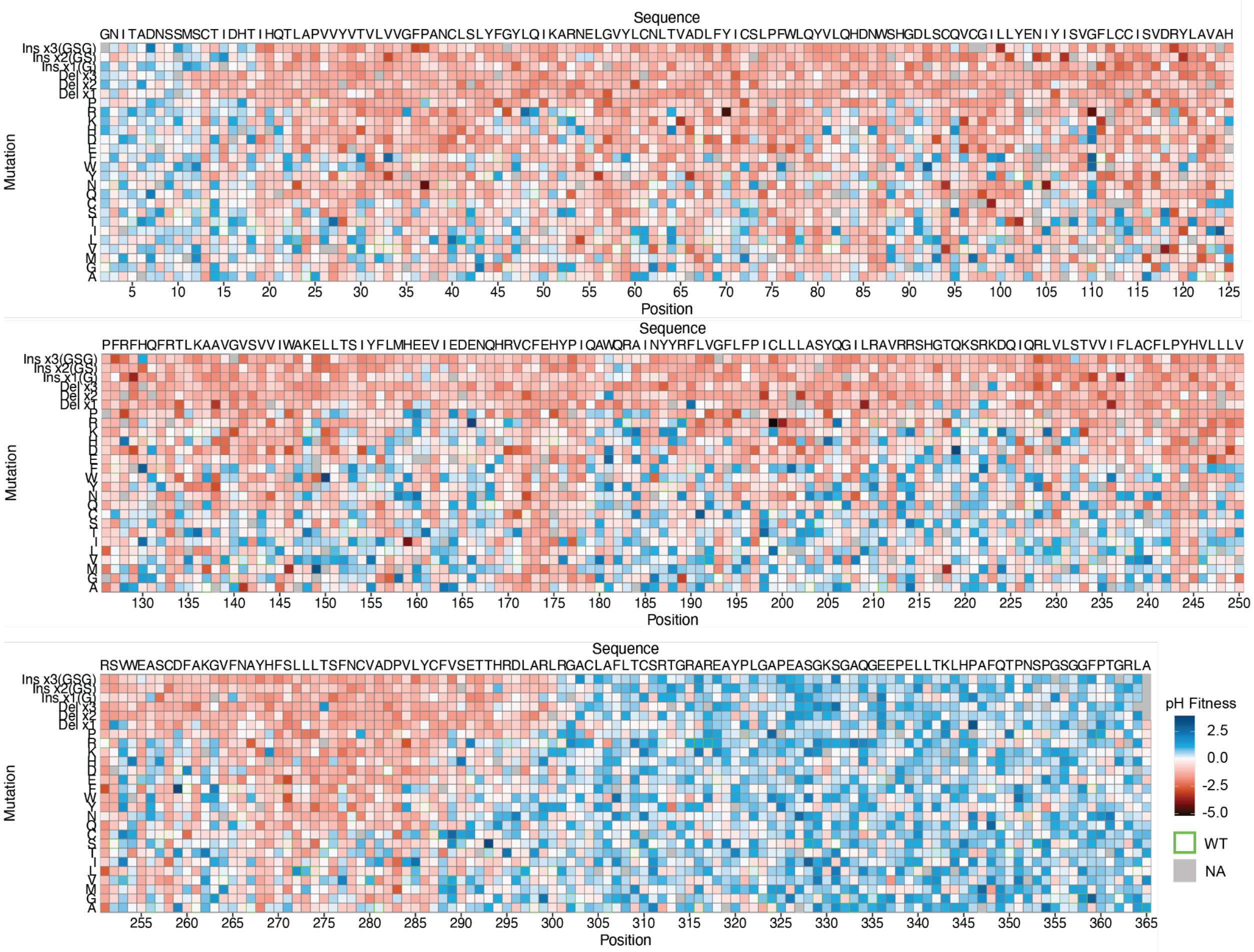


(**A**) Full heatmap of DMS fitness values for GPR68 at pH 5.5. WT sequence is shown above each section of heatmap, mutations are indicated on the left axis of each section, and the amino acid position is indicated by the numbers below each section. Positions and mutations with no data are shown as gray, and the WT amino acid at each position has a green border. Fitness scores are relative to WT and were calculated using Enrich2. Blue indicates increased expression relative to WT, red indicates decreased expression relative to WT. Data are fitness values from three biologically independent deep mutational scans.

### Fig. S10: GPR68 cAMP signaling at pH 5.5 DMS SE heatmap

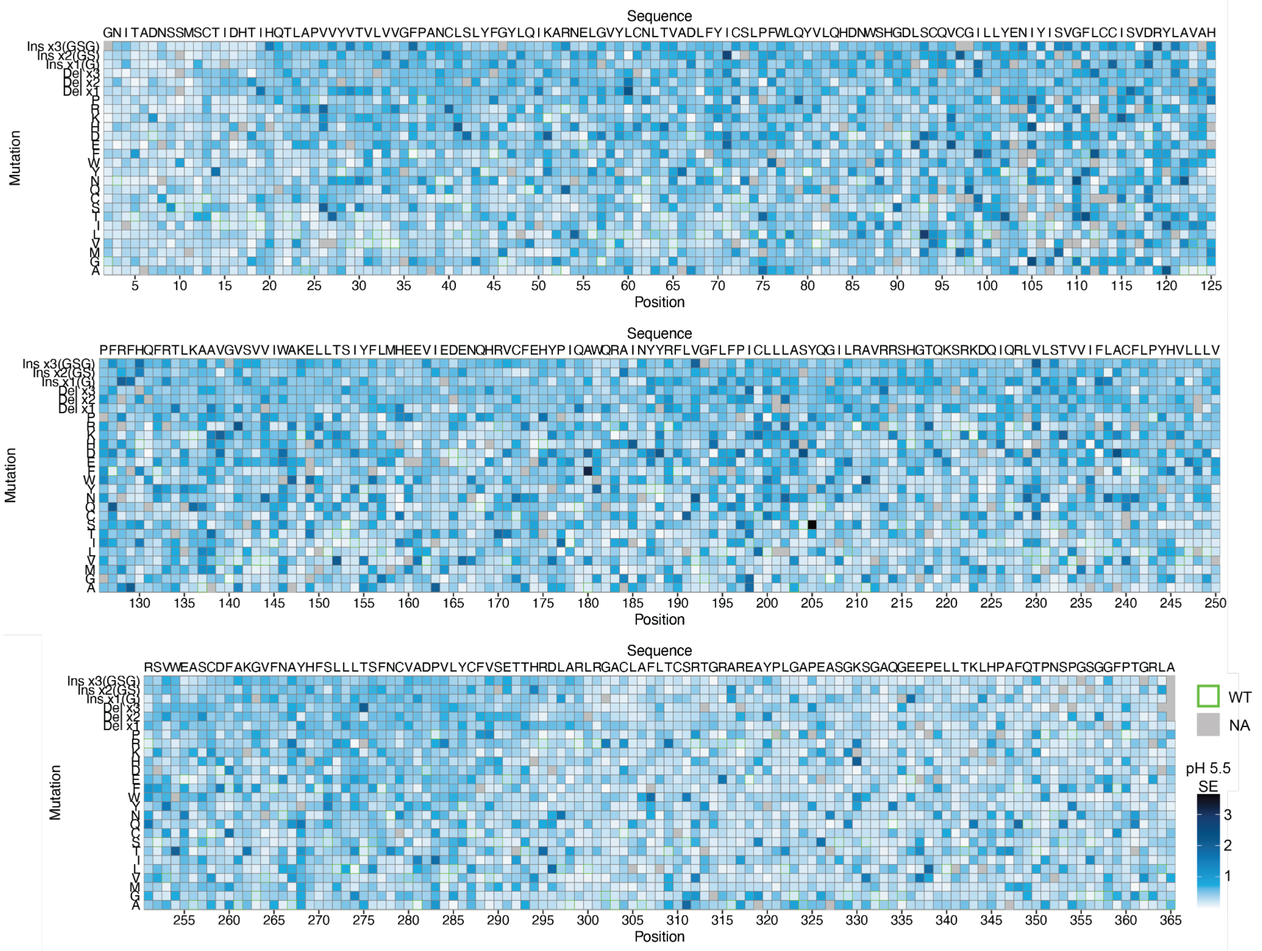


(**A**) Full heatmap of DMS standard errors (SE) for the GPR68 cAMP signaling at pH 5.5 screen. WT sequence is shown above each section of heatmap, mutations are indicated on the left axis of each section, and the amino acid position is indicated by the numbers below each section. Positions and mutations with no data are shown as gray, and the WT amino acid at each position has a green border. SE were calculated using Enrich2 and scaled white to blue. Data are fitness values from three biologically independent deep mutational scans.

**Fig. S11: GPR68 cAMP signaling at pH 6.5 DMS fitness score full heatmap**

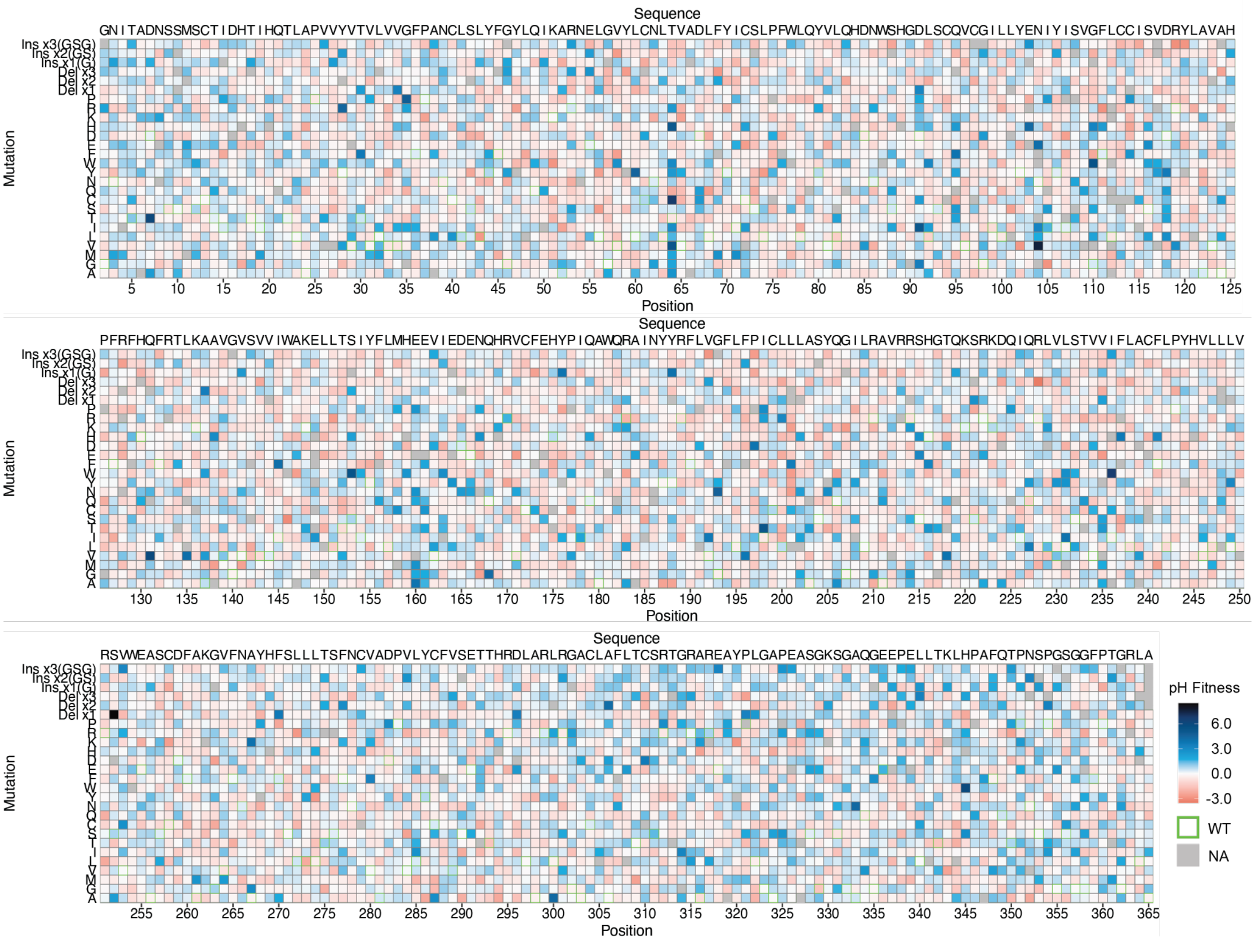


(**A**) Full heatmap of DMS fitness values for GPR68 at pH 6.5. WT sequence is shown above each section of heatmap, mutations are indicated on the left axis of each section, and the amino acid position is indicated by the numbers below each section. Positions and mutations with no data are shown as gray, and the WT amino acid at each position has a green border. Fitness scores are relative to WT and were calculated using Enrich2. Blue indicates increased cAMP signaling relative to WT, red indicates decreased cAMP signaling relative to WT. Data are fitness values from three biologically independent deep mutational scans.

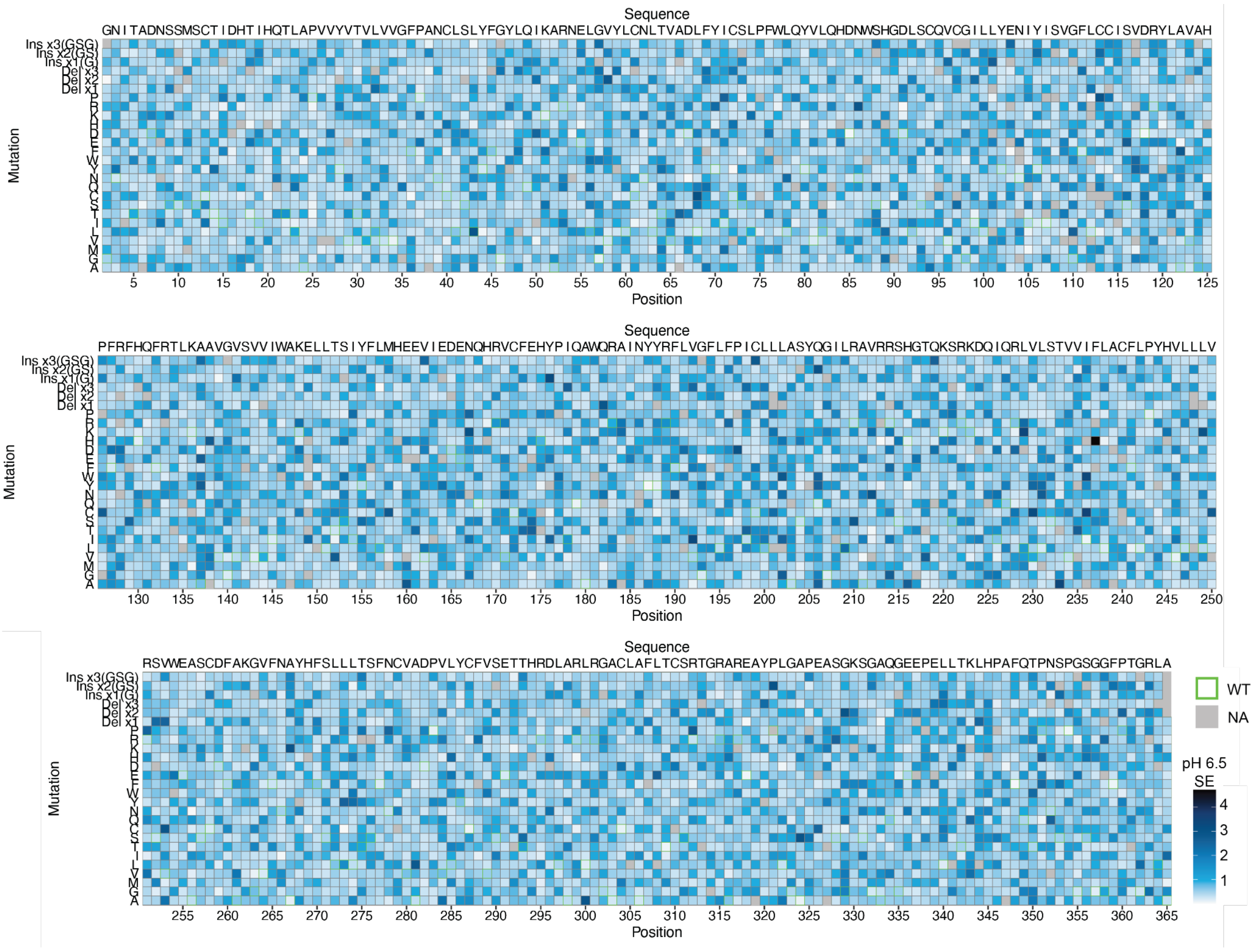


(**A**) Full heatmap of DMS standard errors (SE) for the GPR68 cAMP signaling at pH 6.5 screen. WT sequence is shown above each section of heatmap, mutations are indicated on the left axis of each section, and the amino acid position is indicated by the numbers below each section. Positions and mutations with no data are shown as gray, and the WT amino acid at each position has a green border. SE were calculated using Enrich2 and scaled white to blue. Data are fitness values from three biologically independent deep mutational scans.

**Fig. S13: DMS replicate correlations**

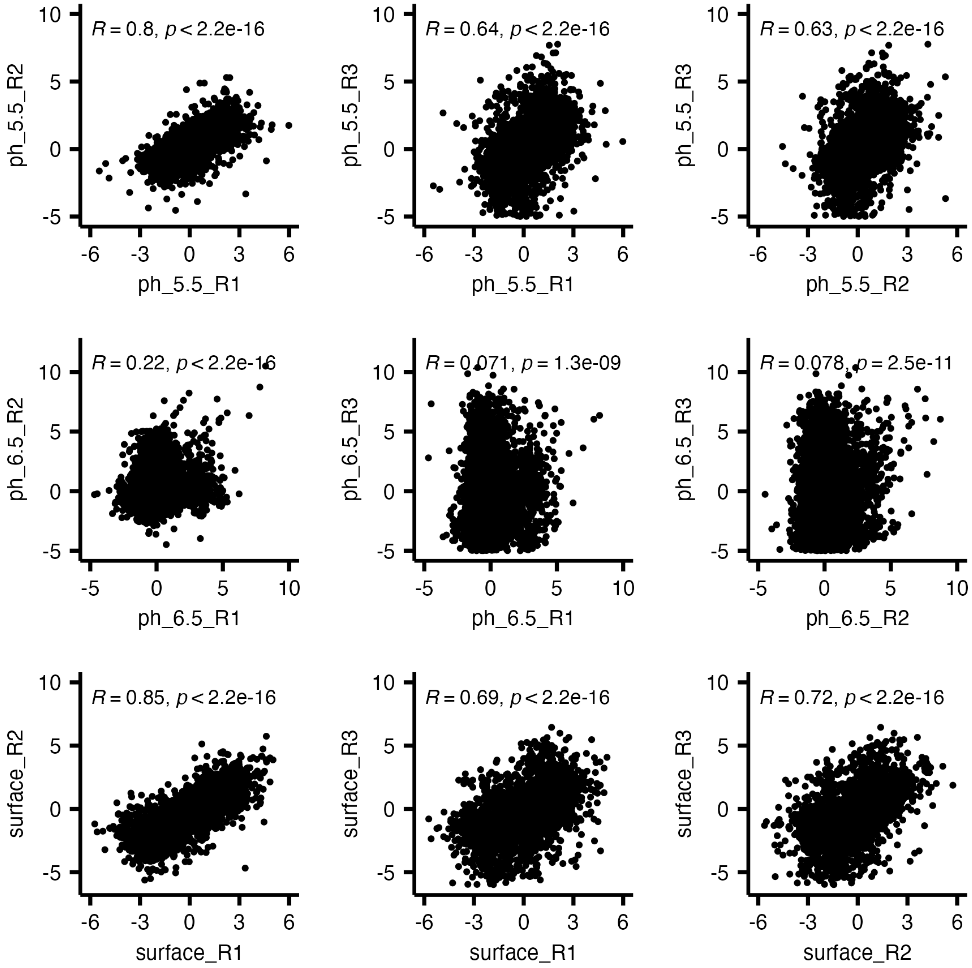


(**A**) Replicate correlations for each DMS screening condition. Spearman coefficient (*R*) and p-values are indicated for each plot.

**Fig. S14: pH 5.5 DMS mean score snapshots**

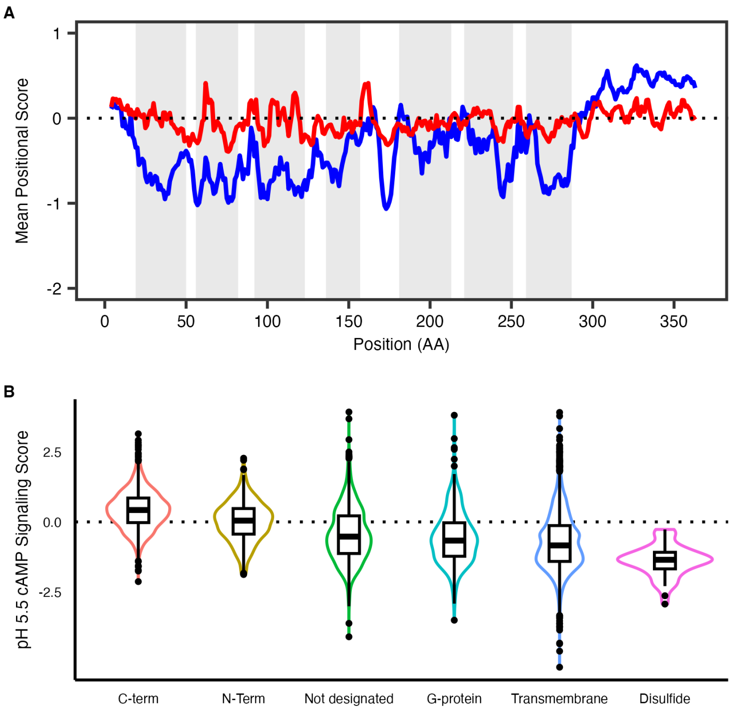


(**A**) Mean missense DMS score at each position along GPR68 for the pH 5.5 (blue) and pH 6.5 (red) screens. Transmembrane helices defined by our structure are indicated in grey. (**B**) Distribution of mean missense mutation scores for GPR68 pH 5.5 DMS for the N-terminus, transmembrane helices (“TM”), C-terminus, disulfides, and Gα contacts (4Å cutoff based on our GPR68-miniG$\alpha_{s/q}$ structure).

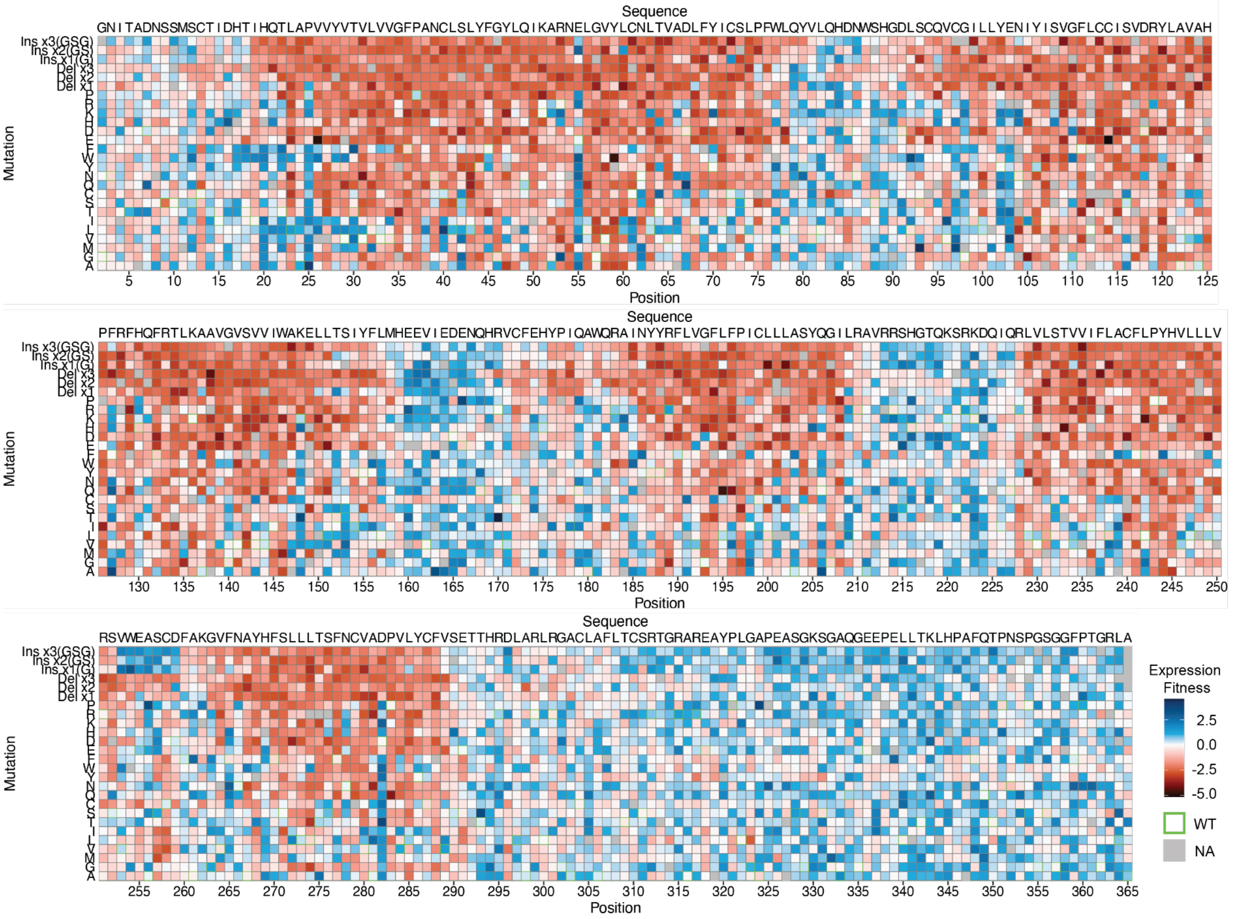


(**A**) Full heatmap of DMS fitness values for GPR68 surface expression. WT sequence is shown above each section of heatmap, mutations are indicated on the left axis of each section, and the amino acid position is indicated by the numbers below each section. Positions and mutations with no data are shown as gray, and the WT amino acid at each position has a green border. Fitness scores are relative to WT and were calculated using Enrich2. Blue indicates increased expression relative to WT, red indicates decreased expression relative to WT. Data are fitness values from three biologically independent deep mutational scans.

### Fig. S16: GPR68 surface expression DMS SE heatmap

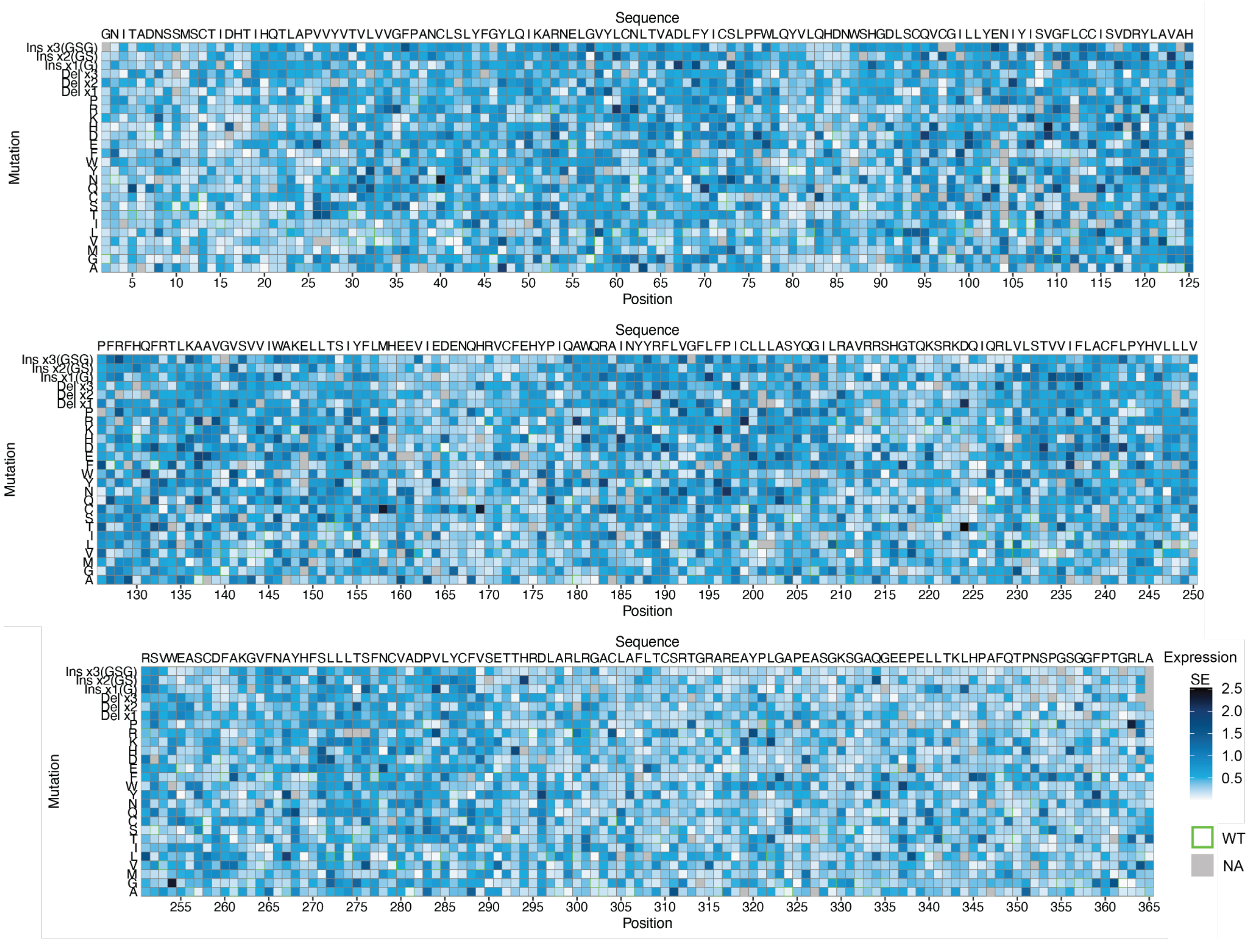


(**A**) Full heatmap of DMS standard errors (SE) for the GPR68 surface expression screen. WT sequence is shown above each section of heatmap, mutations are indicated on the left axis of each section, and the amino acid position is indicated by the numbers below each section. Positions and mutations with no data are shown as gray, and the WT amino acid at each position has a green border. SE were calculated using Enrich2 and scaled white to blue. Data are fitness values from three biologically independent deep mutational scans.

**Fig. S17: Rank plot of surface-adjusted signaling scores from pH 6.5 DMS screen.**

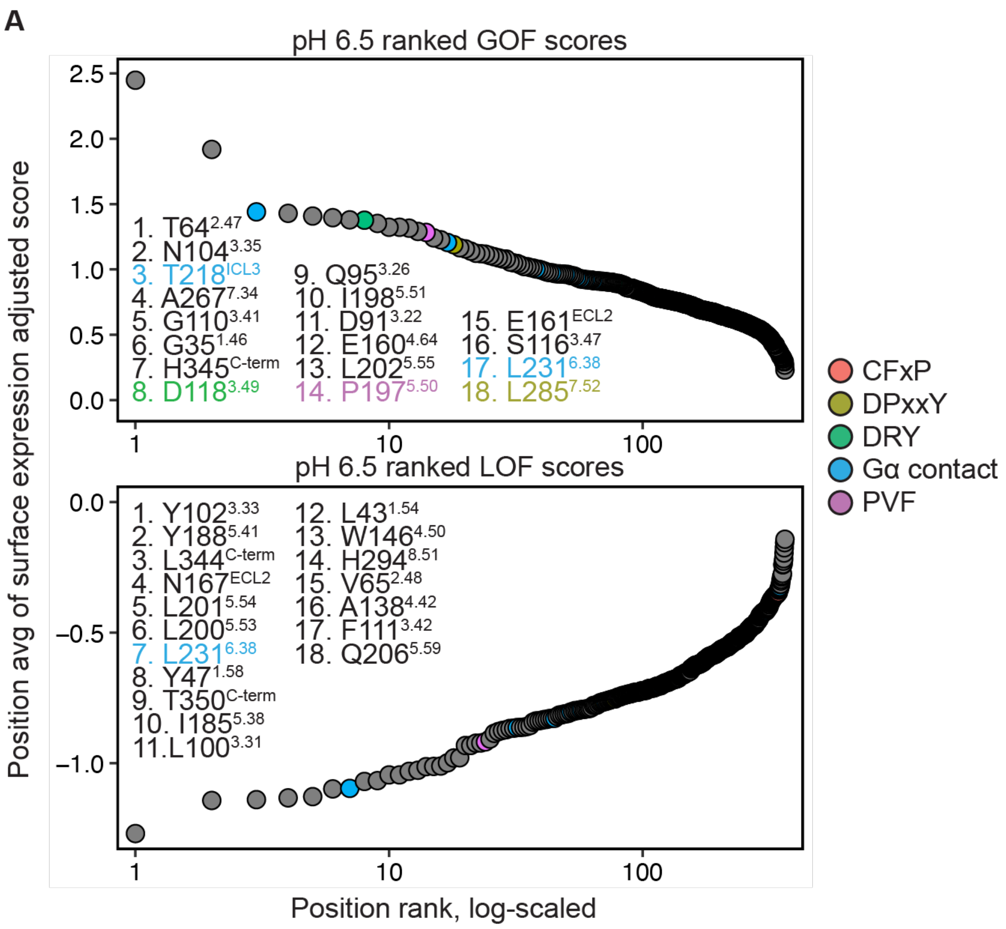


(**A**) Plots of surface expression adjusted cAMP signaling scores from the pH 6.5 deep mutational scan on GPR68. Points are colored as indicated for conserved GPCR sequence motifs. Residues within 4 Å of miniG$\alpha_{s/q}$ in our GPR68 structure are also colored as indicated. The top 5% of positions (LOF/GOF) are annotated within the plot and colored according to sequence motif. Superscripts indicate BW numbering.

**Fig. S18: Additional snapshots of GPR68 GOF and LOF positions.**

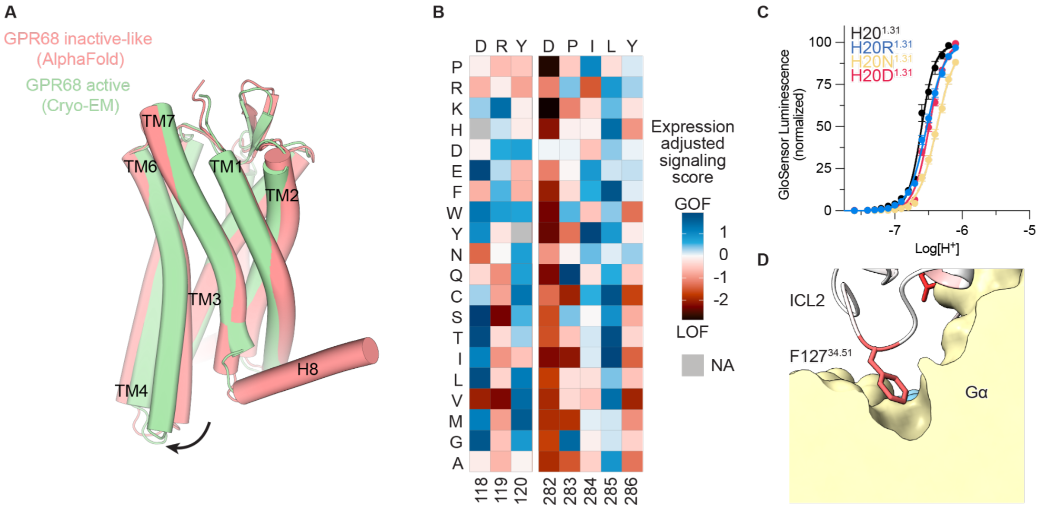


(**A**) Comparison of active-state cryo-EM structure of GPR68 with Alphafold inactive-like structure. (**B**) Heatmap of surface expression-adjusted pH 5.5 cAMP signaling scores for the DRY and DPxxY motifs GPR68. Mutations are indicated on the left axis, and the amino acid position is indicated by the numbers below each. Positions and mutations with no data are shown as gray. Blue indicates higher activity relative to WT, red indicates lower activity relative to WT. (**C**) cAMP accumulation GloSensor assays testing impact of mutations to H20$^{1.31}$ on proton potency in GPR68. Data shown is from three independent biological replicates ± SD. (**D**) Zoom view of GPR68 F127 within ICL2 which is critical for G protein activation. GPR68 residues are colored by LOF score as in **A**.

### Fig. S19: Functional importance of homologous sites in GPR4 and GPR65

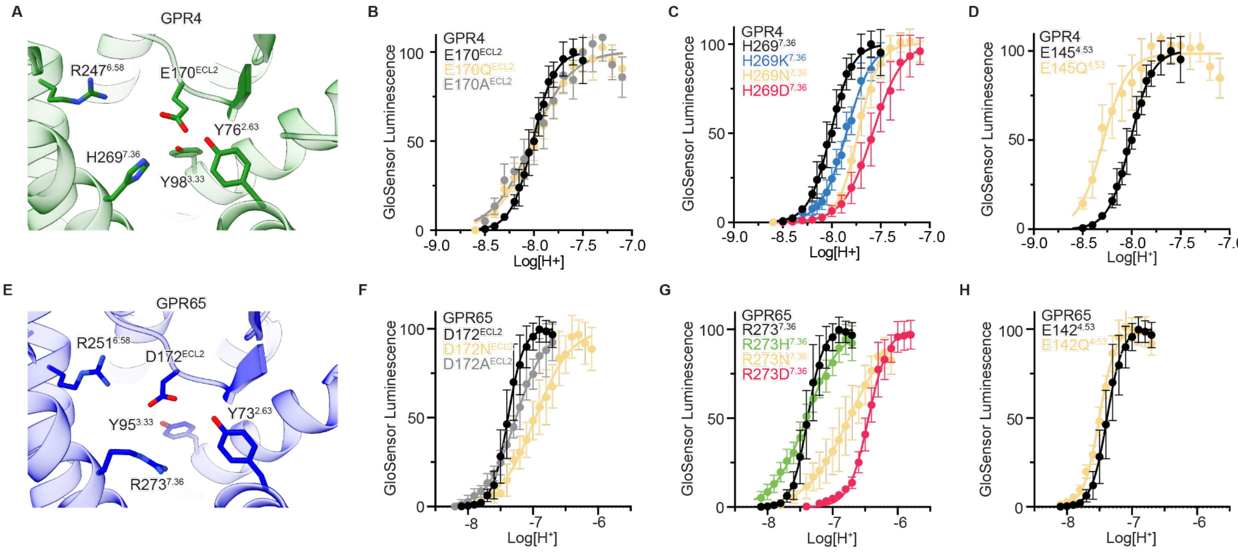


I

```
                                                                  3.33 3.37
GPR4    MGN----HTWEGCHVDSRVDHLFPPSLYIFVIGVGLPTNCLALWAAYRQVQQRNELGVYL  56
GPR65   -------MNSTCIEEQHDLDHYLFPIVYIFVIIVSIPANIGSLCVSFLQAKKESELGIYL  53
GPR68   MGNITADNSSMSCTIDHTIHQTLAPVVYVTVLVVGFPANCLSLYFGYLQIKARNELGVYL  60

GPR4    MNLSIADLLYICTLPLWVDYFLHHDNWIHGPGSCKLFGFIFYTNIYISIAFLCCISVDRY  116
GPR65   FSLSLSDLLYALTLPLWIDYTWNKDNWTFSPALCKGSAFLMYMNFYSSTAFLTCIAVDRY  113
GPR68   CNLTVADLFYICSLPFWLQYVLQHDNWSHGDLSCQVCGILLYENIYISVGFLCCISVDRY  120
                                                 3.34
                            4.53                                45.52
GPR4    LAVAHPLRFARLRRVKTAVAVSSVVWATELGANSAPLFHDELFR-----DRYNHTFCFEK  171
GPR65   LAVVYPLKFFFLRTRRFALMVSLSIWILETIFNAVMLWEDETVVEYCDAEKSNFTLCYDK  173
GPR68   LAVAHPFRFHQFRTLKAAVGVSVVIWAKELLTSIYFLMHEEVIE-----DENQHRVCFEH  175

GPR4    FPMEGWVAWMNLYRVFVGFLFPWALMLLSYRGILRAVRGSVSTERQEKAKIKRLALSLIA  231
GPR65   YPLEKWQINLNLFRTCTGYAIPLVTILICNRKVYQAVRHNKATENKEKKRIIKLLVSITV  233
GPR68   YPIQAWQRAINYYRFLVGFLFPICLLLASYQGILRAVRRSHGTQKSRKDQIQRLVLSTVV  235
                  6.52  6.58                7.36
GPR4    IVLVCFAPYHVLLLSRSAIYLGR--PWDCGFEERVFSAYHSSLAFTSLNCVADPILYCLV  289
GPR65   TFVLCFTPFHVMLLIRCILEHAVNFEDHSNSGKRTYTMYRITVALTSLNCVADPILYCFV  293
GPR68   IFLACFLPYHVLLLVRSVW------EASCDFAKGVFNAYHFSLLLTSFNCVADPVLYCFV  289

GPR4    NEGARSDVAKALHNLLRFLASDKPQEMANASLTLETPLTSKRNSTAKAMTGSWAATPPSQ  349
GPR65   TETGRYDMWNILKFCTGRCNTSQRQRKR--ILSVSTKDTMELEVLE--------------  337
GPR68   SETTHRDLARLRGACLAFLTCSRTGRARE-AYPLGAPEASGKSG--------------AQ  334

GPR4    GDQV--QLKMLPPAQ----------------  362
GPR65   -------------------------------  337     TMs   BW #
GPR68   GEEPELLTKLHPAFQTPNSPGSGGFPTGRLA  365
```

(**A**) Zoom view of residues linking ECL2 of GPR4. (**B**-**D**) GloSensor data for structurally conserved ECL2 residue mutants in GPR4. (**E**) Zoom view of residues linking ECL2 of GPR65. (**F**-**H**) GloSensor data for structurally conserved ECL2 residue mutants in GPR65. Curves in **B-D, F-H** are from three independent biological replicates ± SD. (**I**) Sequence alignment of GPR4, GPR65, and GPR68 with transmembrane helices indicates and BW labels of residues tested in **B-D**, **F-H**, and those that correspond to residues in the GPR68 activation network in **Fig. 6**.

**Table S1: cAMP signaling assay for chimeric GPR4-GPR68 constructs**

| Receptor Chimera | Basal RLU | Emax (Fold) | $pH_{50}$ | Hill |
|---|---|---|---|---|
| GPR4: N-term GPR68 | 289 | 43.2 | 7.92 | 3.07 |
| GPR4: TM2-3 GPR68 | no pH response | | | |
| GPR4: TM4-5 GPR68 | 347 | 7.8 | 7.53 | 2.30 |
| GPR4: TM6-7 GPR68 | 308 | 42.0 | 7.91 | 3.85 |
| GPR4: N-term, TM2-3 GPR68 | no pH response | | | |
| GPR4: N-term, TM4-5 GPR68 | 562 | 4.0 | 7.96 | 2.36 |
| GPR4: N-term, TM6-7 GPR68 | 138 | 16.5 | 7.79 | 2.95 |
| GPR4: TM2-3, TM4-5 GPR68 | no pH response | | | |
| GPR4: TM2-3, TM6-7 GPR68 | no pH response | | | |
| GPR4: TM4-5, TM6-7 GPR68 | 81 | 13.1 | 7.53 | 1.84 |
| GPR4: N-term, TM2-3, TM4-5 GPR68 | no pH response | | | |
| GPR4: N-term, TM2-3, TM6-7 GPR68 | no pH response | | | |
| GPR4: N-term, TM4-5, TM6-7 GPR68 | 332 | 6.0 | 8.03 | 1.89 |
| GPR4: TM2-3, TM4-5, TM6-7 GPR68 | 50 | 3.9 | 7.46 | 2.04 |
| GPR4: N-term,TM2-3, TM4-5, TM6-7 GPR68 | 74 | 33.1 | 7.06 | 1.37 |

**Table S2: Cryo-EM data collection, refinement and validation statistics**

| EMDB: Full map<br>EMDB: 7TM map<br>RCSB PDB: Model | GPR4<br>$G_s$ complex<br>(EMDB-XXXXX)<br>(EMDB-XXXXX)<br>(PDB XXXX) | GPR65<br>$G_s$ complex<br>(EMDB-XXXXX)<br>(EMDB-XXXXX)<br>(PDB XXXX) | GPR68<br>$G_s$ complex<br>(EMDB-XXXXX)<br>(EMDB-XXXXX)<br>(PDB XXXX) | GPR68<br>$G_{s/q}$ complex<br>(EMDB-XXXXX)<br>(EMDB-XXXXX)<br>(PDB XXXX) |
|---|---|---|---|---|
| **Data collection and processing** | | | | |
| Magnification | 105,000 | 105,000 | 105,000 | 105,000 |
| Voltage (kV) | 300 | 300 | 300 | 300 |
| Electron exposure (e–/Å$^2$) | 50 | 45.8 | 49 | 60 |
| Defocus range (μm) | -2.1 to -1.0 | -2.1 to -1.0 | -2.1 to -1.0 | -2.1 to -1.0 |
| Pixel size (Å) | 0.86 (physical) | 0.835 (physical) | 0.83 (physical) | 0.873 (physical) |
| Symmetry imposed | *C1* | *C1* | *C1* | *C1* |
| Initial particle images (no.) | 8,608,607 | 8,673,428 | 7,064,401 | 6,764,523 |
| Final particle images (no.) | 439,296 | 411,592 | 552,937 | 501,967 |
| Map resolution (Å) (masked) | 2.8 | 2.8 | 2.9 | 2.9 |
| FSC threshold | 0.143 | 0.143 | 0.143 | 0.143 |
| **Refinement** | | | | |
| Initial model used (PDB code) | AlphaFold (GPR4)<br>7LJC (G protein)<br>3SN6 (Nb35) | AlphaFold (GPR65)<br>7LJC (G protein)<br>3SN6 (Nb35) | AlphaFold (GPR68)<br>7LJC (G protein)<br>3SN6 (Nb35) | AlphaFold (GPR68)<br>7LJC (G protein)<br>3SN6 (Nb35) |
| Model resolution (Å) (unmasked/masked) | 3.1/3.0 | 4.1/4.0 | 3.4/3.1 | 3.2/3.1 |
| FSC threshold | 0.5 | 0.5 | 0.5 | 0.5 |
| Map sharpening *B* factor (Å$^2$) | -144 | -165 | -133 | -130 |
| Model composition | | | | |
| Non-hydrogen atoms | 8326 | 7716 | 7137 | 7038 |
| Protein residues | 1056 | 969 | 890 | 913 |
| Ligands | 0 | 0 | 1 | 1 |
| *B* factors (Å$^2$) | | | | |
| Protein | 42.75 | 264.60 | 58.86 | 26.24 |
| Ligand | | | 64.97 | 22.64 |
| R.m.s. deviations | | | | |
| Bond lengths (Å) | 0.002 | 0.003 | 0.004 | 0.004 |
| Bond angles (°) | 0.511 | 0.609 | 0.658 | 0.607 |
| Validation | | | | |
| MolProbity score | 1.53 | 1.72 | 2.49 | 1.84 |
| Clashscore | 2.73 | 8.77 | 14.65 | 4.59 |
| Poor rotamers (%) | 1.91 | 0.12 | 0 | 1.95 |
| CaBLAM outliers (%) | 1.16 | 1.48 | 0.40 | 1.15 |
| Ramachandran plot | | | | |
| Favored (%) | 96.26 | 96.25 | 98.12 | 94.39 |
| Allowed (%) | 3.74 | 3.75 | 1.88 | 5.61 |
| Disallowed (%) | 0 | 0 | 0 | 0 |

**Table S3: cAMP signaling assay for GPR68 mutants**

| Receptor | Basal RLU ± SEM | Emax (fold) ± SEM | $pH_{50}$ ± SEM | Hill ± SEM |
|---|---|---|---|---|
| WT | 95.2 ± 1.2 | 46.0 ± 1.3 | 6.66 ± 0.00 | 4.16 ± 0.07 |
| H20D | 83.3 ± 1.9 | 46.6 ± 1.5 | 6.52 ± 0.00 | 3.43 ± 0.04 |
| H20N | 77.2 ± 2.8 | 43.5 ± 2.3 | 6.45 ± 0.02 | 2.96 ± 0.03 |
| H20R | 80.8 ± 0.8 | 40.0 ± 1.0 | 6.55 ± 0.01 | 3.29 ± 0.09 |
| E149Q | 186.6 ± 74.7 | 27.4 ± 2.6 | 7.41 ± 0.05 | 2.60 ± 0.32 |
| E174A | 102.0 ± 1.3 | 23.6 ± 1.3 | 6.73 ± 0.01 | 2.76 ± 0.10 |
| E174Q | 93.3 ± 4.2 | 18.0 ± 1.2 | 6.16 ± 0.01 | 2.69± 0.05 |
| R251A | 113.6 ± 5.2 | 23.0 ± 1.6 | 5.84 ± 0.01 | 2.88 ± 0.12 |
| R251D | 116.4 ± 7.0 | 17.1 ± 1.6 | 5.77 ± 0.00 | 3.06 ± 0.01 |
| H269D | 104.3 ± 3.1 | 44.3 ± 1.5 | 6.34 ± 0.01 | 2.94 ± 0.06 |
| H269K | 115.9 ± 58.8 | 28.2 ± 7.4 | 6.87 ± 0.03 | 2.59 ± 0.11 |
| H269N | 99.4 ± 3.0 | 53.5 ± 3.0 | 6.46 ± 0.01 | 2.71 ± 0.09 |

**Table S4: cAMP signaling assay for GPR4 mutants**

| Receptor | Basal RLU ± SEM | Emax (Fold) ± SEM | $pH_{50}$ ± SEM | Hill ± SEM |
|---|---|---|---|---|
| GPR4 WT | 213.3 ± 20.5 | 39.1 ± 3.7 | 8.01 ± 0.01 | 4.20 ± 0.24 |
| GPR4 H16D | 364.3 ± 114.9 | 40.4 ± 5.6 | 7.80 ± 0.01 | 3.32 ± 0.23 |
| GPR4 H16N | 303.1 ± 113.1 | 51.7 ± 6.6 | 7.85 ± 0.01 | 3.88 ± 0.20 |
| GPR4 H16R | 204.1 ± 47.5 | 65.8 ± 10.4 | 7.75 ± 0.01 | 3.34 ± 0.15 |
| GPR4 E145Q | 943.8 ± 83.6 | 5.7 ± 0.3 | 8.27 ± 0.02 | 3.65 ± 0.35 |
| GPR4 E170A | 582.7 ± 78.5 | 8.2 ± 1.0 | 8.02 ± 0.04 | 2.24 ± 0.08 |
| GPR4 E170Q | 522.3 ± 66.7 | 16.4 ± 1.4 | 7.98 ± 0.00 | 2.28 ± 0.16 |
| GPR4 R247A | 246.1 ± 29.4 | 21.9 ± 1.9 | 7.81 ± 0.03 | 2.53 ± 0.07 |
| GPR4 R247D | 241.6 ± 66.9 | 19.7 ± 7.3 | 7.49 ± 0.01 | 1.81 ± 0.17 |
| GPR4 H269D | 127.0 ± 20.6 | 117.4 ± 21.4 | 7.55 ± 0.03 | 3.01 ± 0.17 |
| GPR4 H269N | 122.6 ± 12.5 | 111.2 ± 11.0 | 7.72 ± 0.01 | 4.21 ± 0.17 |
| GPR4 H269K | 181.2 ± 9.1 | 60.2 ± 3.4 | 7.85 ± 0.01 | 3.35 ± 0.12 |

**Table S5: cAMP signaling assay for GPR65 mutants**

| Receptor | Basal RLU ± SEM | Emax (fold) ± SEM | $pH_{50}$ ± SEM | Hill ± SEM |
|---|---|---|---|---|
| GPR65 | 109.3 ± 11.2 | 79.9 ± 10.5 | 7.39 ± 0.03 | 4.31 ± 0.28 |
| GPR65 H13D | 180.2 ± 20.0 | 71.4 ± 4.7 | 7.12 ± 0.03 | 2.10 ± 0.08 |
| GPR65 H13N | 198.8 ± 17.4 | 40.6 ± 3.7 | 7.11 ± 0.03 | 1.94 ± 0.10 |
| GPR65 H13R | 149.4 ± 10.1 | 46.1 ± 3.7 | 7.04 ± 0.03 | 2.22 ± 0.09 |
| GPR65 Y95F | 169.5 ± 12.2 | 57.1 ± 5.1 | 7.18 ± 0.02 | 3.37 ± 0.13 |
| GPR65 E142Q | 337.0 ± 44.4 | 25.0 ± 2.8 | 7.50 ± 0.01 | 3.68 ± 0.13 |
| GPR65 D172A | 149.0 ± 33.7 | 10.0 ± 2.5 | 7.25 ± 0.02 | 2.08 ± 0.11 |
| GPR65 D172N | 666.7 ± 299.8 | 15.6 ± 3.2 | 6.99 ± 0.03 | 1.71 ± 0.07 |
| GPR65 R249A | 162.9 ± 22.5 | 39.8 ± 18.3 | 6.57 ± 0.02 | 4.88 ± 0.19 |
| GPR65 R249D | 86.4 ± 13.7 | 13.9 ± 3.9 | 6.97 ± 0.10 | 1.23 ± 0.07 |
| GPR65 R273D | 210.4 ± 24.7 | 29.6 ± 4.9 | 6.46 ± 0.03 | 3.08 ± 0.13 |
| GPR65 R273H | 1031.2 ± 142.7 | 5.6 ± 0.5 | 7.40 ± 0.04 | 1.72 ± 0.05 |
| GPR65 R273N | 438.9 ± 84.8 | 16.7 ± 3.6 | 6.80 ± 0.06 | 1.69 ± 0.09 |
